# Monte Carlo Wavelet Analysis for Objective Peak Detection in SRM LC-MS/MS Analysis

**DOI:** 10.64898/2025.12.18.694988

**Authors:** Randall K. Julian, Brian A. Rappold, Fan Yi, Stephen R. Master

## Abstract

Detection of low-level analytes in complex chromatographic-mass spectrometric data requires a criterion to discern apparent peaks from background. Conventional signal-to-noise criteria rely on simple, constant-variance noise models and overlook spurious peaks generated by chemical noise and co-eluting interferences. We introduce a wavelet-based Monte Carlo technique for determining the statistical significance of SRM LC-MS/MS peaks in the presence of structured chemical noise. The method empirically characterizes chemical-noise peaks in samples and builds a generative noise-only null model. Monte Carlo resampling of the noise model assigns *p*-values that are controlled for the family-wise type I error rate (FWER). We validated the method with SRMs from a dilution series of drug compounds in plasma with known ground-truth concentrations. Triplicate technical replicates were used, spanning concentrations from far above the limit of detection to far below it. Peaks with adjusted *p <* 0.05 matched the expectation for true positives above the detection limit. Peaks below the limit of detection matched matrix blanks as true negatives, and intermittent detection in the transition region was observed. An independent external validation using a clinical pain panel confirmed the method detects ketamine in confirmed positive samples with signal intensity below the lowest calibration standard while correctly classifying matrix blanks and biological negatives. As a demonstration, we applied our method to a recently published lipid mediator data set. By replacing subjective noise-region selection with a formal hypothesis test against an empirical null model, the method provides an objective and reproducible criterion for deciding whether peak integration is warranted.

---

Liquid chromatography-tandem mass spectrometry (LC-MS/MS) in selected reaction monitoring (SRM) mode is a powerful method for quantifying low-abundance biological molecules in complex samples. At the concentration levels where many SRM measurements occur, the dominant analytical challenge is not instrument sensitivity but instead structured chemical noise that has analyte-like characteristics^1,2^. Systematic studies of atmospheric-pressure ionization LC-MS have identified ionic chemical background noise as a significant limitation in trace analysis, with families of recurring background ions and reproducible precursor-product patterns that survive into MS/MS spectra^35^. In MALDI mass spectrometry, chemical-noise background is likewise recognized as the main factor limiting practical sensitivity, because decreasing analyte load eventually yields signals indistinguishable from chemical noise^6^.

These issues raise particular challenges for characterizing putative analyte peaks using criteria such as the signal-to-noise (S/N) ratio. Standard S/N thresholds are typically justified for electronic noise models, yet at trace levels, the dominant source of false positives is chemical clutter. A fixed S/N cutoff (e.g., 3 or 5) requires an analyst to choose where noise is measured. The selection of a region free from clutter is subjective and makes the S/N value for any given peak ambiguous. In practice, apparent peaks are often integrated to yield areas that are converted to concentrations. Integrating chemical noise peaks produces spurious concentration estimates that propagate unchecked through the analysis. This applies equally to relative abundance comparisons: a chemical noise peak integrated at the expected retention time contributes a spurious area that distorts fold-change estimates and differential abundance statistics regardless of whether an absolute concentration is ever calculated. Related peak-quality approaches have used raw-EIC shape and residual-based quality metrics for targeted and untargeted LC-MS data, but they do not provide a formal hypothesis test against a chemical noise null model with chromatogram-wide error control over false-positive rates^7,8^.

To address this challenge, we introduce and validate a statistical approach that estimates and simulates chemical noise in order to assess whether apparent peaks represent genuine signals. In targeted SRM, the data for a given transition and matrix are one-dimensional time series, *I*(*t*), measured as a function of retention time *t*. Chromatographic peaks can be approximated by a Gaussian shape above 50% of their height but span a wide range of widths and amplitudes. We accommodate this variability using a continuous wavelet transform (CWT) with a Mexican hat (DOG2, Marr) mother wavelet to identify peaks across wavelet scales in blank and sample chromatograms.

By excluding the narrow retention-time window around the internal-standard-aligned analyte peak, we empirically characterize the chemical-noise process: distributions of peak locations, effective widths (via wavelet scale), and powers (wavelet energy). These components define the null model for 1-D SRM chromatograms, in which simulated data contain only baseline noise plus a realistic number of randomly located chemical-noise peaks drawn from empirical distributions built directly from study samples.

To assess the likelihood that a feature can be explained by the null model, we adopt Hopes simplified Monte Carlo significance test^9^. We generate a reference set of random chromatograms under the noise-only null hypothesis, evaluate wavelet power at each time point and wavelet scale, and compute *p*-values by counting how often the null power exceeds the observed power. This approach allows hypothesis testing for complex null models even when they cannot be expressed in closed form.

Because this procedure is performed at every time point and wavelet scale, the resulting matrix of *p*-values represents a multiple-testing problem. We collapse the multiscale wavelet information to a single statistic at each time point (the minimum *p*-value across wavelet scales) and control the family-wise error rate (FWER) across all time points using the Holm-Bonferroni step-down procedure^10^. The FWER is the probability of obtaining at least one false-positive detection anywhere in the chromatogram. This differs fundamentally from simple S/N thresholds, which treat each decision as isolated and offer no chromatogram-wide error guarantee. We define analyte presence as rejection of the chemical noise null model (FWER-adjusted p < 0.05) at the expected retention time in the quantifier SRM transition, and absence as failure to reject it.

Previous applications of wavelet analysis to chromatographic^11^ and mass-spectral data^12^ have largely focused on peak picking and denoising under electronic and proportional noise models. More broadly, as Maraun and Kurths emphasized, naive use of theoretical background spectra in wavelet significance testing can misrepresent significance in the presence of correlated noise and multiple testing across wavelet scales and times^13^. In contrast, the present approach replaces idealized noise models with an empirically derived, generative model of chemically structured noise, couples it to multiscale wavelet analysis, and provides explicit control of family-wise error across the chromatogram.

This approach is related to constant false alarm rate (CFAR) detection in radar signal processing^14^. In CFAR, detection thresholds are adapted to local clutter statistics (the local background fluctuation pattern) to maintain a constant false-alarm probability as the background varies. Rather than estimating clutter from a sliding window, we use a global model derived from sample (and matrix blank) chromatograms to simulate the distribution of wavelet power under the null hypothesis. The goal is analogous: to set thresholds that control false positives with respect to realistically modeled clutter, rather than idealized noise baselines.

We validate the Monte Carlo approach using a dilution series of drugs of known concentration spiked into plasma. We also show the application of this method using publicly available data from the RvT4 study of Walker et al. (2024)^15^ a targeted SRM dataset as a test case involving a different compound and different instrumentation. To help with making the approach as usable as possible, all the source code is available from the repository https://github.com/rkjulian/wavelet-peak-significance.

## Methods

### Data Acquisition

#### Validation Dilution Series

Certified reference material of clonidine (1.0 mg/mL, Supelco, C-033), gabapentin (1.0 mg/mL, Supelco, G-007S), and lorazepam (1.0 mg/mL, Supelco, L-901) were purchased from MilliporeSigma (St. Louis, MO, USA). Stable isotope-labeled internal standards (IS), including clonidine-D_4_ (0.1 mg/mL, Supelco, C-157), gabapentin-^13^C_3_ (0.1 mg/mL, Supelco, G-018), and lorazepam-D_4_ (0.1 mg/mL, Supelco, L-902), were also obtained from Millipore-Sigma. Methanol (Optima LC/MS Grade, A456-500), acetonitrile (ACN) (Optima LC/MS Grade, A955-4), isopropanol (IPA) (Optima Grade, A464-4) and formic acid (FA) (Optima LC/MS Grade, A117-50) were purchased from Fisher Scientific (Waltham, MA, USA). Ultrapure water (Milli-Q grade, 18.2 MΩ·cm) was produced using a Milli-Q IQ 7000 system (Millipore, Burlington, MA, USA). Human K_2_EDTA plasma (Lot# HMN1475449) was purchased from BioIVT (Westbury, NY, USA) and served as the biological matrix.

A mixed analyte working solution (1 *µ*g/mL for each analyte) was prepared by combining appropriate volumes of each reference standard solution and diluting to 1 mL with methanol. A 5 ng/mL spiked plasma sample was prepared by diluting 2 *µ*L of the analyte working solution into 400 *µ*L of blank plasma.

Serial two-fold dilutions of the 5 ng/mL spiked plasma sample with blank human plasma were prepared to generate a concentration range from 0.0195 to 5.00 ng/mL for each of the three analytes. Dilution steps resulted in the following final nominal concentrations: 0.0195, 0.0391, 0.0781, 0.156, 0.313, 0.625, 1.25, 2.50, and 5.00 ng/mL. These samples are named 0.25X, 0.5X, 1X, 2X, 4X, 8X, 16X, 32X, 64X (See Table 2).

A mixed IS working solution (1 *µ*g/mL for each IS) was prepared by combining 2 *µ*L of each labeled internal standard and diluting to 200 *µ*L with methanol. The IS working solution was further diluted 20-fold to 50 ng/mL in 50/50 (v/v) methanol/water to prepare the IS spike solution.

#### Sample Preparation

Reagent blank (water), plasma blank, and spiked plasma samples (20 *µ*L each, prepared in triplicate) were combined with 10 *µ*L of IS spike solution and mixed at 1000 rpm for 1 min. Protein precipitation was performed by adding 200 *µ*L ACN, followed by vortexing at 1000 rpm for 5 min at room temperature. Samples were centrifuged at 2500 × *g* for 5 min, and 50 *µ*L of the supernatant was transferred and evaporated under nitrogen. Residues were reconstituted in 200 *µ*L of 90/10 (v/v) water/ACN, mixed for 1 min at 1000 rpm, and transferred to glass vials for LC-MS/MS analysis.

#### LC-MS/MS Conditions

The extracted samples were analyzed using a Waters Acquity I-Class Plus UPLC system coupled with a Xevo TQ Absolute XR triple quadrupole mass spectrometer (Waters, Milford, MA, USA) with a 5 *µ*L injection volume.

A Waters Acquity UPLC BEH C_18_ column (50 × 2.1 mm, 1.7 *µ*m; SKU 186002350) was used for chromatographic separation. Mobile phase (MP) A consisted of water with 0.1% FA, and MP B consisted of ACN with 0.1% FA. The gradient was programmed as follows: 4% MP B from 0 to 0.5 min; linear increase to 95% B from 0.5 to 3.0 min; held at 95% B from 3.0 to 4.0 min; and re-equilibrated at 4% B from 4.01 to 5.0 min. The flow rate was 0.5 mL/min. The weak needle wash and seal wash consisted of 90:10 (v/v) water/ACN, while the strong needle wash consisted of 2:2:2:2:0.1 (v/v) water/ACN/IPA/MeOH/FA.

Analytes and IS were monitored using multiple reaction monitoring (MRM) in positive electrospray ionization (ESI) mode. The MRM transitions, dwell time, cone voltages, and collision energies for quantifier ions are summarized in Table 1.

**Table 1:** Quantifier MRM transitions and stable-isotope-labeled internal standards for the validation dilution-series compounds. All transitions were monitored in positive ESI mode on the Waters Xevo TQ Absolute XR with a dwell time of 0.017 s and cone voltage of 30 V.

| Compound | Q1 (m/z) | Q3 (m/z) | CE (V) |
| --- | --- | --- | --- |
| Gabapentin | 172.3 | 154.3 | 13 |
| Gabapentin-$^{13}\text{C}_3$ (IS) | 175.3 | 157.4 | 9 |
| Clonidine | 232.2 | 215.1 | 23 |
| Clonidine-$\text{D}_4$ (IS) | 236.2 | 219.0 | 23 |
| Lorazepam | 321.2 | 275 | 35 |
| Lorazepam-$\text{D}_4$ (IS) | 325.2 | 233.0 | 35 |

**Table 2:**
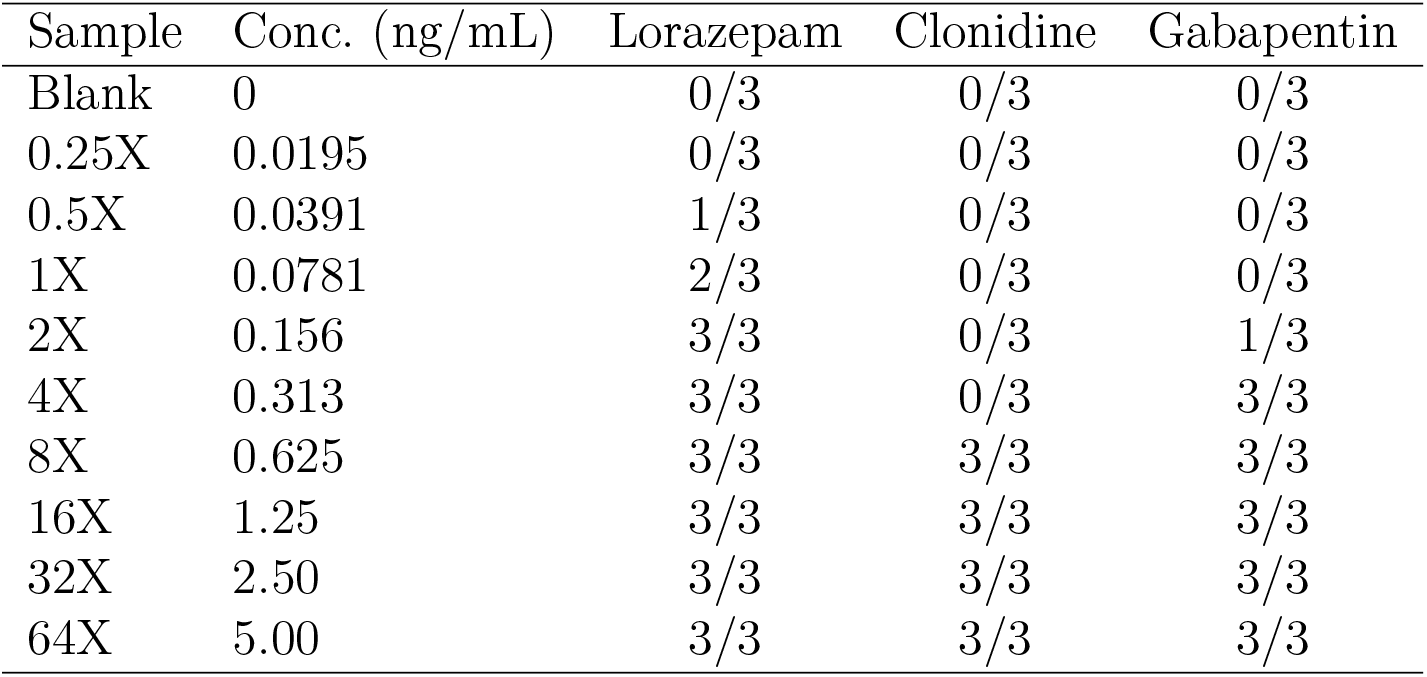
Wavelet-based detection results for the drug dilution series in K_2_EDTA plasma (Waters Xevo TQ Absolute XR). Each cell shows the number of replicates detected out of 3. The 64X concentration (5.00 ng/mL) served as the positive-control standard.

For all samples in the dilution series analyses, both noise characterization and hypothesis testing were restricted to a 0.6-3.0 min analysis window (600 scans). This window encompasses the full analyte elution range for all three compounds while ensuring that the stationary-noise assumptions underlying the null model are satisfied.

#### Retention-Time Alignment

For each analyte we defined the expected retention time in every sample by aligning to the internal standard in the highest concentration sample. We first measured the offset between the analyte and IS peaks in a standard or high concentration sample chromatogram, ΔRT = RT_analyte,std_ −RT_IS, std_, and then, for each lower concentration sample, we set the expected analyte retention time to RT_analyte_ = RT_IS, sample_ + ΔRT. In all figures, the IS retention time is shown as a solid vertical line and the IS-aligned analyte retention time as a black dashed vertical line.

### Software

All chromatogram processing, wavelet analyses, and Monte Carlo simulations were performed in R version 4.5.2^16^ using the MSnbase package for mzML import and chromatogram handling^1719^. The MassSpecWavelet package was used for continuous wavelet transforms^20,21^. The NADA^22^package was used to estimate electronic noise (mean and standard deviation) for the dilution series data, and the truncnorm package^23^ for sampling from truncated normal distributions to add the electronic noise component of the null model. Data wrangling, parallelisation, and figure generation used standard R packages for table manipulation, parallel computing, and plotting. All source code used is available from the repository https://github.com/rkjulian/wavelet-peak-significance.

### Continuous Wavelet Transform Analysis

To detect transient chromatographic features, we treated each SRM chromatogram as a time series and applied the CWT following the approach of Torrence and Compo^24^. In their work, wavelet power at a given time and scale can be compared against a noise background to obtain a local test for significance. The DOG2 wavelet acts as a matched filter for Gaussian-shaped chromatographic peaks, producing maximum response when the wavelet scale matches the peak width. Wavelet power is then compared to the noise model as described in the Implementation section.

#### Mathematical Framework

We compute the CWT *W* (*s, b*) of each chromatogram using the DOG2 mother wavelet^24^. We define wavelet power by squaring positive CWT coefficients and setting negative coefficients to zero:

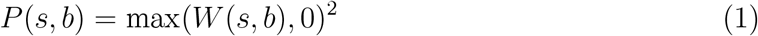

This truncation prevents negative coefficients from producing artifacts in the power map after squaring.

#### Implementation

For each chromatogram, we computed the CWT using the DOG2 wavelet as implemented in the MassSpecWavelet R package^21^ (wavelet = mexh). Wavelet scales were spaced linearly from *s* = 1 to *s*_max_ = min ((*N*− 1)*/*16, 64, where *N* is the number of time points. The upper bound enforces a Nyquist-like constraint: at least two sampling intervals per effective wavelet width, given that the DOG2 kernel spans [− 8, 8]. For hypothesis testing we sampled 100 scales across this range. This conservatively fine grid ensures that no chromatographic peak width falls between adjacent scales. Fewer scales might suffice, but we traded longer computation for reduced risk of false negatives. The chemical-noise characterization uses a narrower 31-scale grid restricted to *s* ∈ [1.5, 3.0] (see Algorithm Summary below). The Monte Carlo testing procedure (Methods, Statistical Testing and Control of False Positive Peaks) operates on the resulting multiscale power fields to assign *p*-values.

### Monte Carlo Null Hypothesis Generation

Torrence and Compo^24^ derived analytical chi-square distributions for wavelet power under white and red noise assumptions. However, chromatographic chemical noise, consisting of discrete interference peaks with non-parametric properties, lacks an analytical representation. This limitation raises concerns about mis-specified background spectra and inflated significance that have been noted for wavelet applications in other fields^13^. Because chromatographic chemical noise lacks such an analytical distribution, we estimate significance by Monte Carlo simulation under the empirically characterized null model (described below). Because this comparison is repeated at many time points and wavelet scales, we then apply a multiple-testing correction^9,10,25^ to control the FWER.

For each Monte Carlo iteration *j* = 1 to *N*_MC_, we construct a simulated noise-only chromatogram by combining three components: chemical noise, unknown interference, and instrumental noise. Each component is described in the following subsections.

### Chemical Noise Component

For each compound and transition in the dilution series, we first summarised the number of recurring chemical-noise peaks per chromatogram from the chemical-noise characterisation (Results, Chemical Noise Characterization). With *n* = 30 chromatograms per compound per transition (blanks through the highest concentration level), the empirical peak-count distributions were well sampled, so we drew the number of recurring chemical-noise peaks *N*_exist_ in each simulated chromatogram from a Poisson distribution with rate *λ* equal to the observed mean peak count for that compound and transition, capped at the number of available observed retention times.

For the gabapentin quantifier transition (m/z 172.3 → 154.3), the chromatograms also contain a second, later-eluting compound that is chromatographically resolved from gabapentin but shares the same SRM channel. To prevent either genuine compound from influencing the chemical-noise characterization, we applied separate exclusion windows around the internal-standard-aligned retention times of both gabapentin and the co-monitored compound, removing all peaks within ±0.03 min of each expected RT before constructing the empirical noise distributions. The co-monitored compound therefore serves as an internal consistency check on the wavelet-based detection (it yields a strong, significant peak at its own RT), but was not analysed quantitatively in this study. For clonidine and lorazepam, whose quantifier transition channel contain only a single target analyte, a single exclusion window around the analyte retention time sufficed.

We sampled the locations (time) and wavelet scales of the *N*_exist_ peaks from the distributions estimated from the chemical-noise characterisation for that transition, and sampled their powers from a kernel density estimate of the observed peak powers. We then added *N*_new_ = 1 additional chemical-noise peak to represent a previously unobserved interference. The rationale is as follows: if the signal observed at the expected retention time were due to chemical noise rather than the target analyte, at least one contaminant peak not accounted for by the recurring chemical-noise model must have appeared at that retention time. Because this contaminant is unknown, its retention time is drawn uniformly from the chromatographic window rather than from the observed location distribution. We set *N*_new_ to 1, the minimum number of additional interfering peaks required to produce a false signal at the target retention time. This keeps the null model as permissive as possible and avoids an overly conservative test that could inflate false negatives. The wavelet scale and power of this additional peak are drawn from the same KDEs as the recurring peaks, so that the interference resembles observed chemical noise in shape and intensity.

Each peak contributes a Gaussian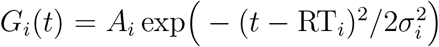, where 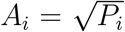, *σ*_*i*_ is proportional to *s*_*i*_ ·Δ*t* (wavelet scale converted to time units), and the constant of proportionality is chosen so that simulated peaks have similar widths to observed peaks at the same wavelet scale.

### Electronic Noise Component

Electronic noise *ϵ*(*t*) is sampled independently at each time point from a truncated normal distribution TruncatedNormal(*µ*_blank_, *σ*_blank_, lower = 0). Because a substantial fraction of reagent-blank intensities fall at or below zero (the detectors censoring threshold), naive sample statistics are biased: censored zeros inflate the mean and compress the variance. For example, 43% of pooled reagent-blank intensities for the lorazepam quantifier transition (m/z 321.2 → 275) were non-positive, yielding naive estimates of 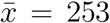 and *s* = 312. To account for this left-censoring, we estimated *µ*_blank_ and *σ*_blank_ using regression on order statistics (ROS)^26^, as implemented in the NADA R package^22^. ROS was applied without transformation, pooling intensities from three replicate reagent-blank injections across the analysis time window (0.6-3.0 min) for each SRM transition. For the lorazepam quantifier, this yielded 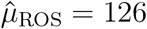 and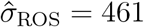.

#### Complete Noise-Only (Null) Chromatogram Model

We define each simulated noise-only chromatogram (our null model) as

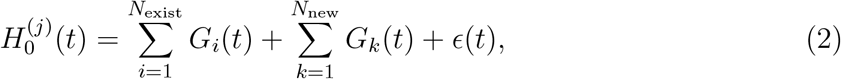

We then apply the CWT to each 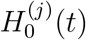 and compute its wavelet power spectrum.

#### Algorithm Summary

The chemical-noise characterization procedure is formalized in Algorithm 1. The algorithm applies wavelet analysis to all chromatograms, identifies peaks outside the analyte window, merges detections across wavelet scales, and constructs distributions of peak properties that define the null model.

For the chemical-noise peak characterization, we focused on wavelet scales corresponding to typical chromatographic peak widths and excluded the smallest, noise-dominated scales. We computed the DOG2 CWT on a restricted scale band *s* ∈ [1.5, 3.0] with step size 0.05 in the dimensionless scale parameter, and detected peaks only within this range (Algorithm 1). The resulting distributions of peak scale and power were then used to parameterise the null model, which itself is evaluated across the full multiscale range described in the Implementation section. We represent these distributions using kernel density estimates (KDEs).

To locate local power maxima, the wavelet power at each scale was lightly smoothed with a 3-point moving average to suppress single-point power fluctuations before wavelet-power peak picking. We defined the prominence of each candidate power-domain peak as the height of the peak above the higher of the adjacent local minima when walking left and right until a higher point or the trace boundary was reached, and retained only power peaks with prominence greater than a fixed absolute threshold *P*_min_ = 1 in wavelet-power units. This near-zero floor avoids degenerate zero-weight artifacts during cross-scale merging; the substantive filtering is performed by the cross-scale consistency and spike-width criteria described below.

After merging detections across scales, two filters removed spurious candidates. First, candidates whose half-height width in the raw chromatogram was fewer than *w*_min_ = 3 data points were discarded as electronic-noise spikes, which are characteristically narrow (1-2 points). Second, candidates had to appear at a minimum of *n*_min_ = 5 of the 31 wavelet scales analyzed (≈16% of the scale band). Real chromatographic peaks produce wavelet responses that persist across a range of scales, whereas noise produces responses at only one or two^21,27^. Together, these filters retained peaks of varying chromatographic width while rejecting isolated electronic noise spikes.

For each chromatogram, we treated a narrow window around the internal-standard-aligned analyte retention time as a protected region and excluded peaks in this region from the chemical-noise characterization. Specifically, we removed all peaks within ±*δ*_RT_ of the aligned analyte retention time before constructing the distributions of chemical-noise peak locations (time), wavelet scales, and powers (Algorithm 1). This notch ensures that any true analyte signal, if present, does not contaminate the noise model. In all analyses, we used *δ*_RT_ = 0.03 min (analyte exclusion half-window) and *δ*_merge_ = 0.05 min (cross-scale merge tolerance).

##### Algorithm 1

Chemical Noise Characterization

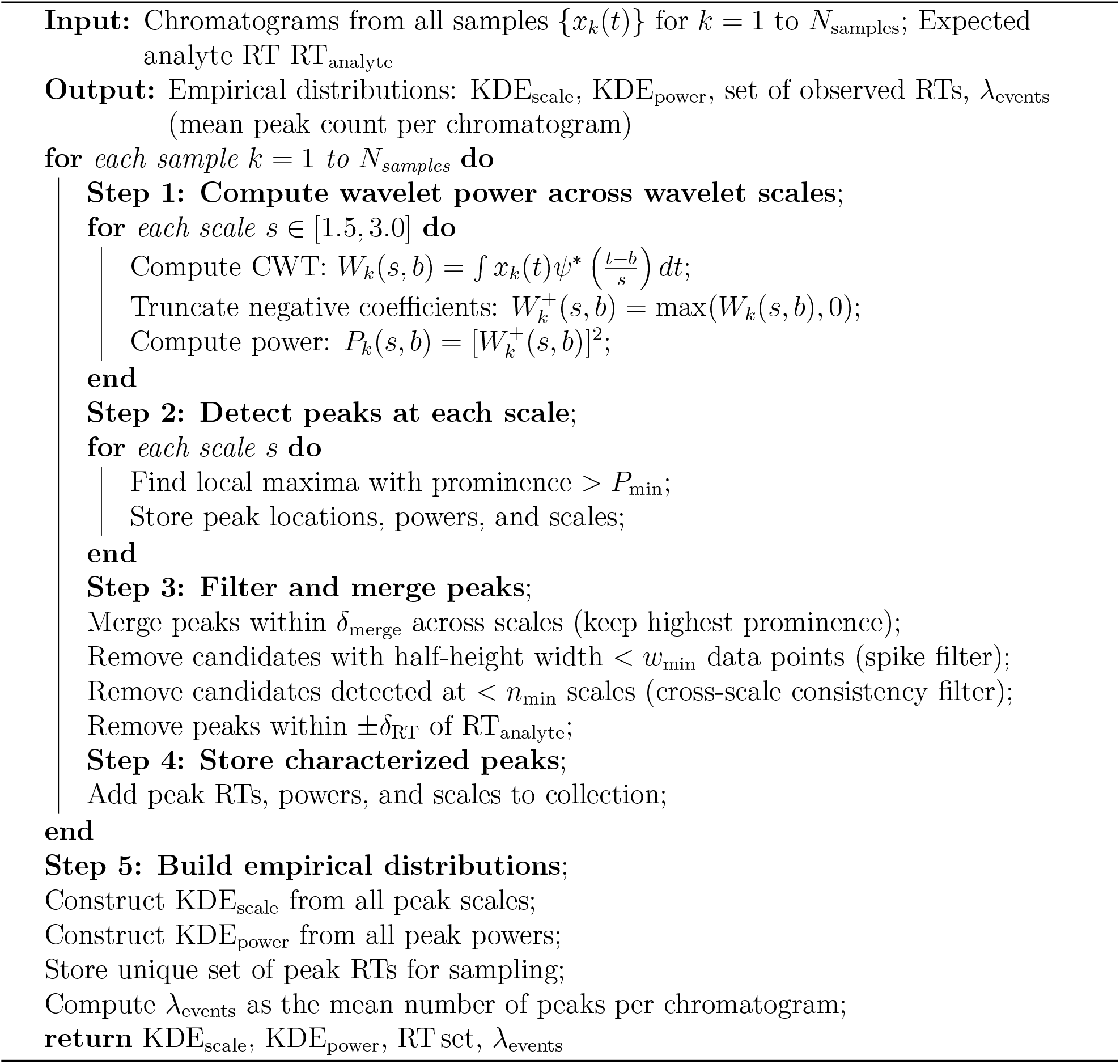

### Statistical Testing and Control of False Positive Peaks

For each time point *b* and wavelet scale *s*, we compare the observed wavelet power to the null distribution and compute *p*(*s, b*) as the fraction of Monte Carlo simulations whose noise-only power equals or exceeds the observed power (plus one, following Hopes correction^9^). We then take the minimum *p*-value across all wavelet scales at each time point, *p*_min_(*b*) = min_*s*_ *p*(*s, b*), so that peaks of any width can be detected provided they exceed the chemical-noise background at some scale. Finally, we apply the Holm-Bonferroni step-down procedure^10^ to the vector {*p*_min_(*b*)} to control the FWER, and declare time points with *p*_adj_(*b*) *< α* = 0.05 as statistically significant. The complete procedure is formalized in Algorithm 2.

## Results

### Chemical Noise Characterization

Using the wavelet-based peak-finding procedure described in Methods, we quantified chemical-noise peaks in all matrix containing chromatograms. For each transition, we counted distinct wavelet peaks outside the analyte retention-time window and summarized their locations, wavelet scales, and powers. Figure 1 illustrates this procedure for a single lorazepam chromatogram.

**Figure 1.**
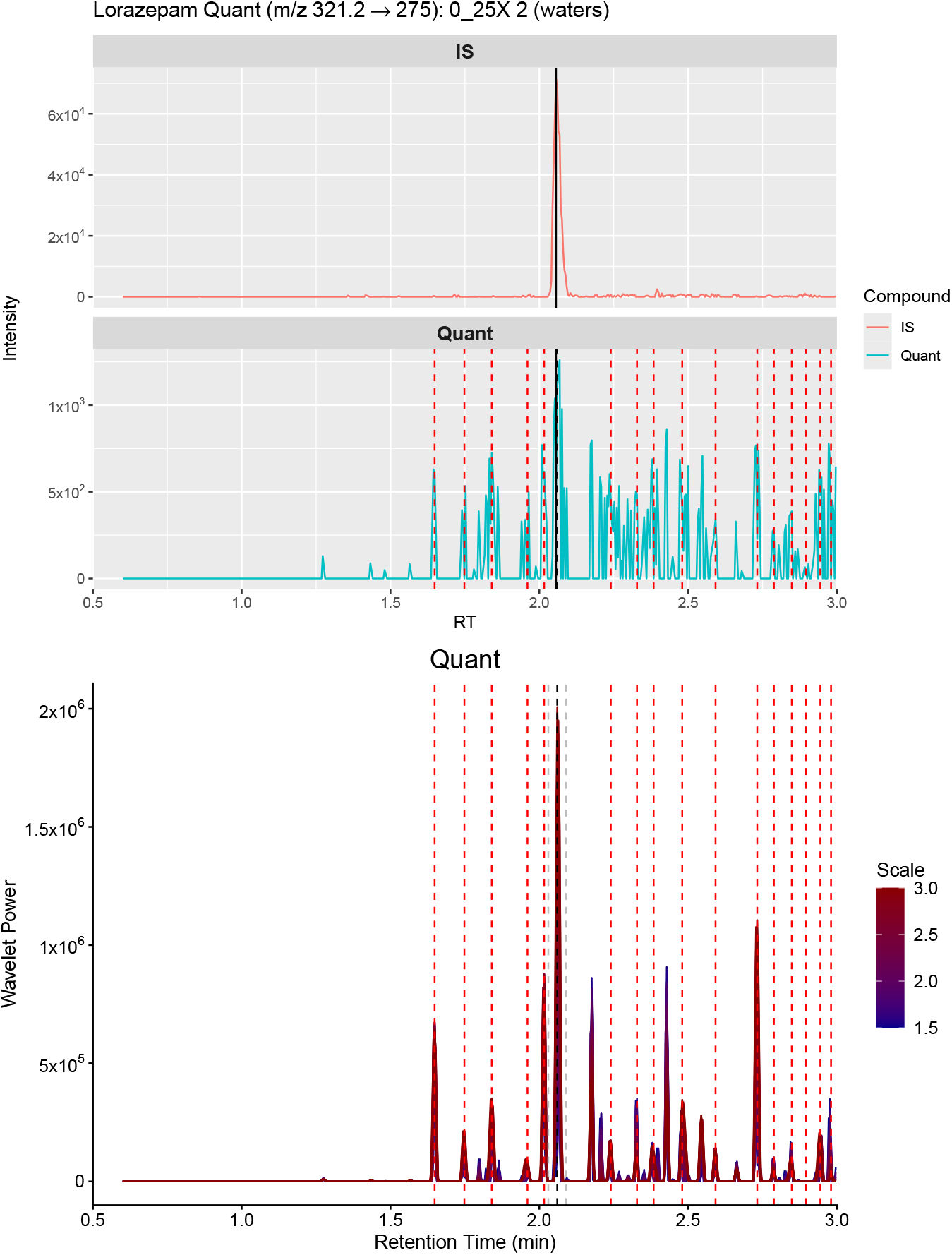
Wavelet-based identification of chemical-noise peaks for a single lorazepam chromatogram (Waters, 0.25X). Top: internal-standard (IS) and quantifier (Quant: m/z 321.2 → 275) chromatograms. Bottom: DOG2 wavelet power across time and scale, with detected chemical-noise peaks marked (red dashed lines). Grey dashed lines indicate the ±0.03 min exclusion window around the internal-standard-aligned lorazepam retention time.

As described in Methods, the number of recurring chemical-noise peaks per simulated chromatogram is drawn from a Poisson distribution with rate *λ*_events_ equal to the observed mean peak count for each compound and transition (*n* = 30 chromatograms each). For the Waters drug dilution data, the DOG2 wavelet analysis yielded *λ*_events_ = 17.3 for the lorazepam quantifier channel (m/z 321.2 → 275), *λ*_events_ = 47.0 for the clonidine quantifier channel (m/z 232.2 → 215.1), and *λ*_events_ = 45.9 for the gabapentin quantifier channel (m/z 172.3 → 154.3).

In the gabapentin quantifier transition channel, a second, later-eluting peak is present and chromatographically separated from gabapentin. The internal-standard-aligned gabapentin and second-analyte retention-time regions were both excluded from the chemical-noise characterization, so that neither true analyte peak contributes to the noise distributions.

The empirical distributions of chemical-noise peak times and powers illustrate two important features (Figure 2). First, noise peaks are mostly uniform at later retention times, but have some structure early in the trace. Second, the distribution of peak powers is broad, with a long right tail of relatively intense noise peaks whose wavelet powers approach those of low-level analyte signals. These observations motivate a null model that explicitly includes discrete chemical-noise peaks rather than treating baseline noise as purely homoscedastic and Gaussian.

**Figure 2.**
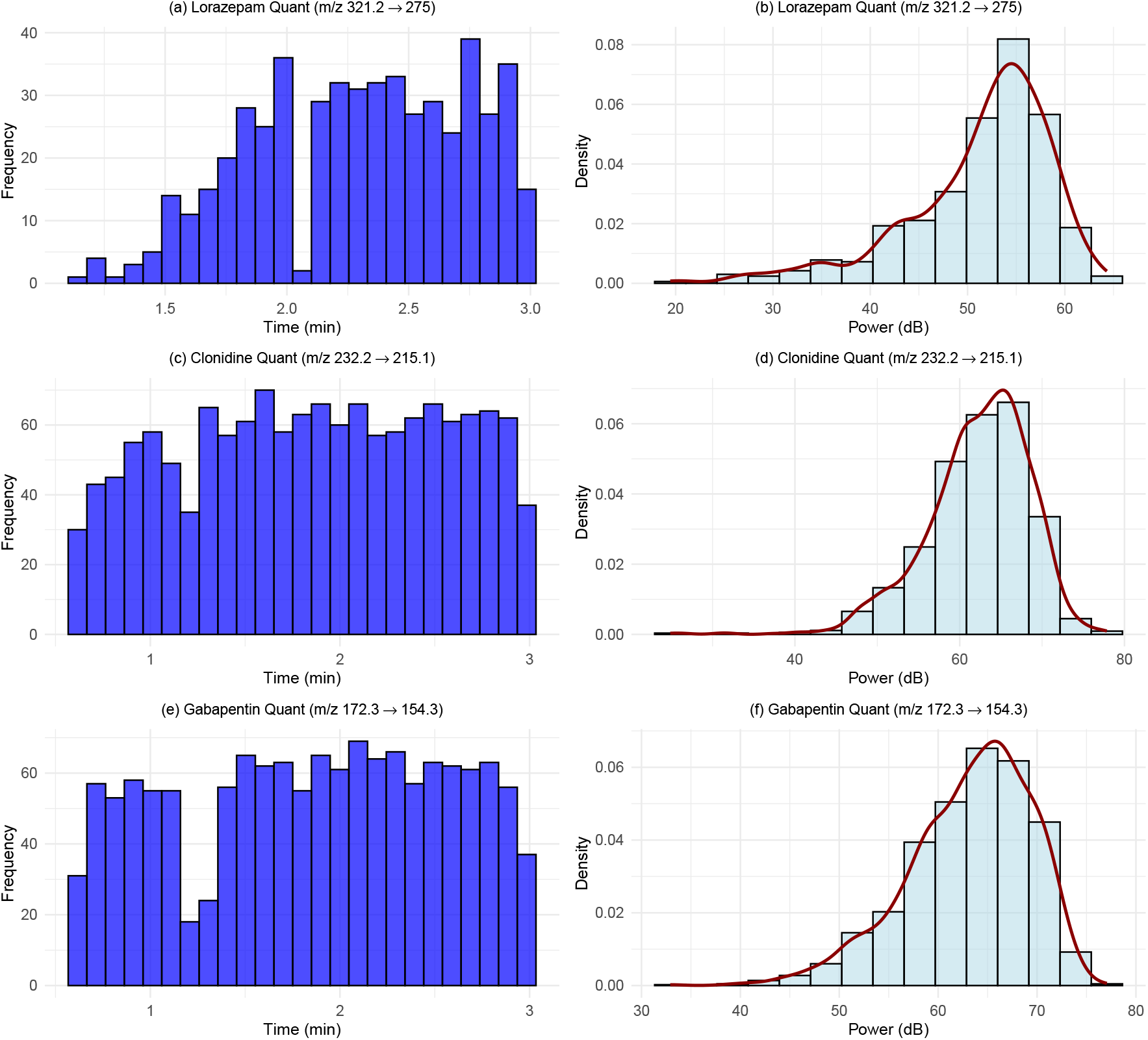
Distributions of chemical-noise peak retention times (left column) and wavelet powers in dB (right column) for the lorazepam (a, b; m/z 321.2 → 275), clonidine (c, d; m/z 232.2 → 215.1), and gabapentin (e, f; m/z 172.3 →154.3) quantifier transitions on the Waters instrument.

### Simulated Noise-Only Chromatograms

To demonstrate the behavior of the null model, we generated example noise-only chromatograms following the procedure described in Methods (Monte Carlo Null Hypothesis Generation). Example null chromatograms for the lorazepam quantifier transition (Figure 3) show discrete but randomly located peaks with widths and amplitudes comparable to those observed in plasma samples, confirming that the generative model produces realistic chromatographic structure.

**Figure 3.**
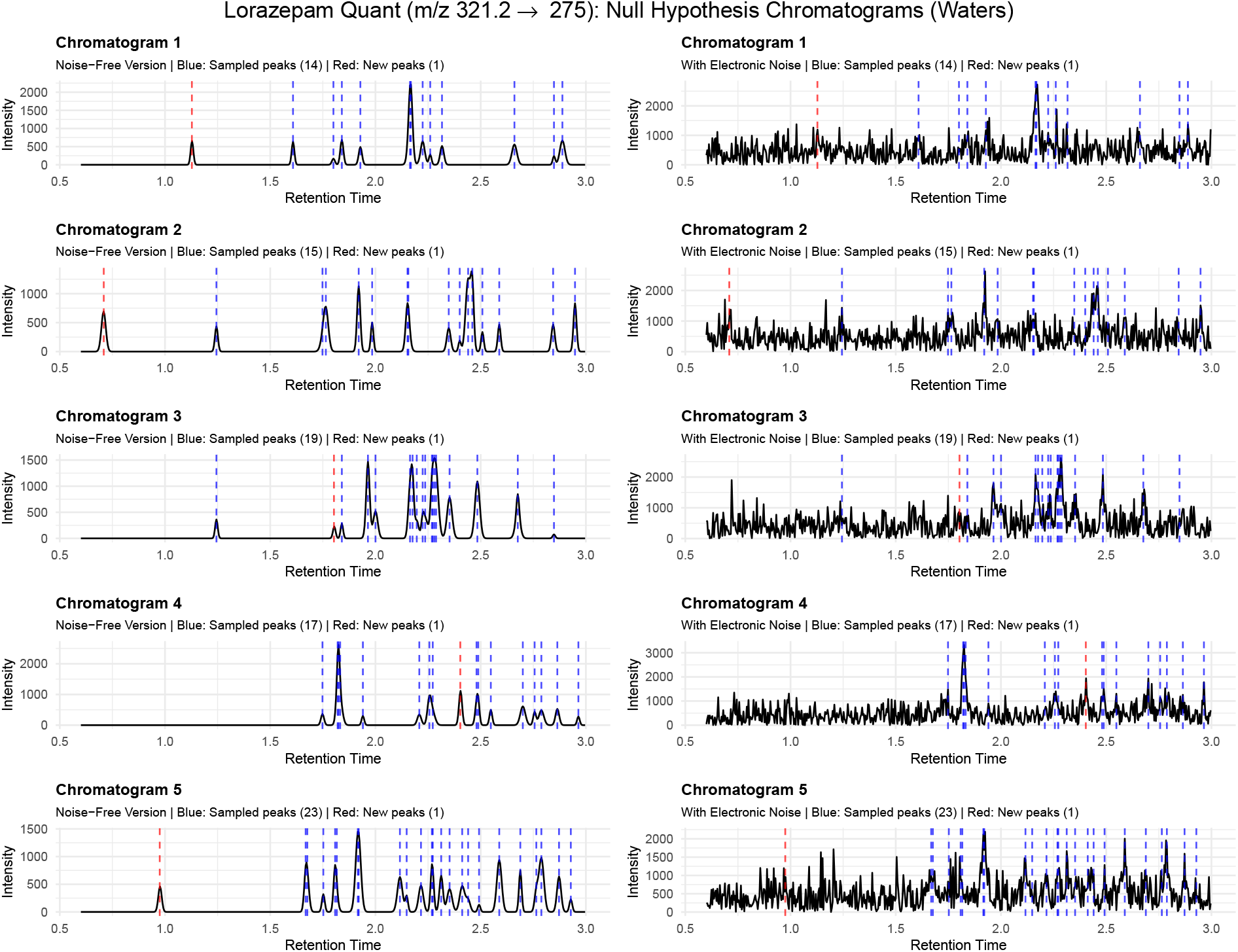
Representative null-hypothesis chromatograms for the lorazepam quantifier channel (m/z 321.2 → 275) on the Waters instrument. Left: chemical-interference peaks only (blue dashed: sampled from observed locations; red dashed: uniformly placed). Right: complete null chromatograms with electronic noise added.

#### Algorithm 2

Wavelet-Based Hypothesis Testing

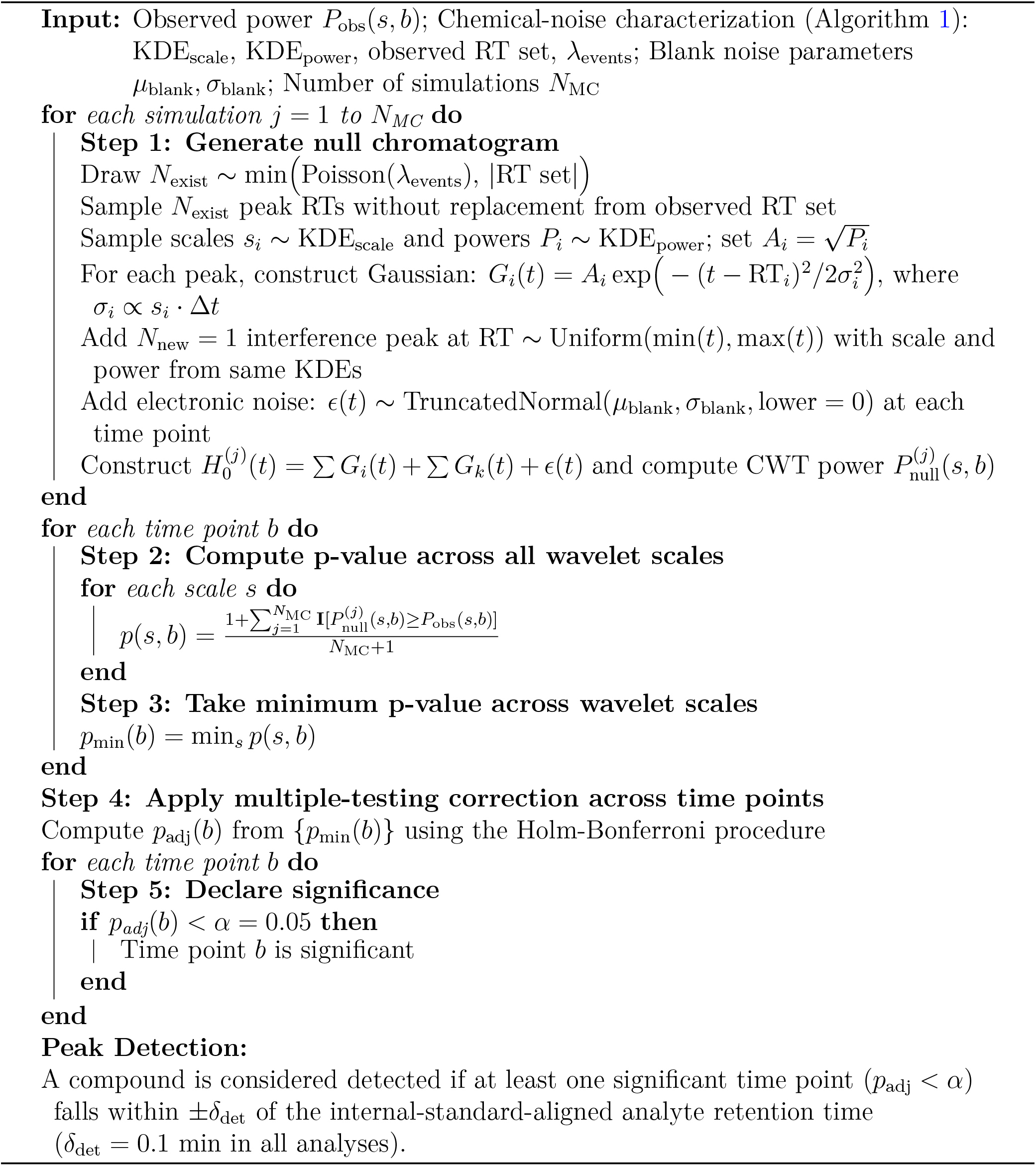

### Validation with Drug Dilution Series

To validate the wavelet-based test, we applied it to the dilution-series data for all three compounds (Lorazepam, Clonidine, and Gabapentin) acquired on the Waters Xevo TQ Absolute XR (Table 1). We compared blank plasma samples, which should contain no analyte signal, against the highest-concentration standard (64X, 5.00 ng/mL), which serves as a positive control for each compound.

Figure 4 compares blank and 64X standard chromatograms with their wavelet-based significance maps for each compound. Note that all wavelet power heatmaps are scaled using an asinh function to account for the large dynamic range in LC-MS/MS chromatograms; the heatmap intensities therefore give a qualitative sense of the wavelet power distributions across wavelet scales and retention time.

**Figure 4.**
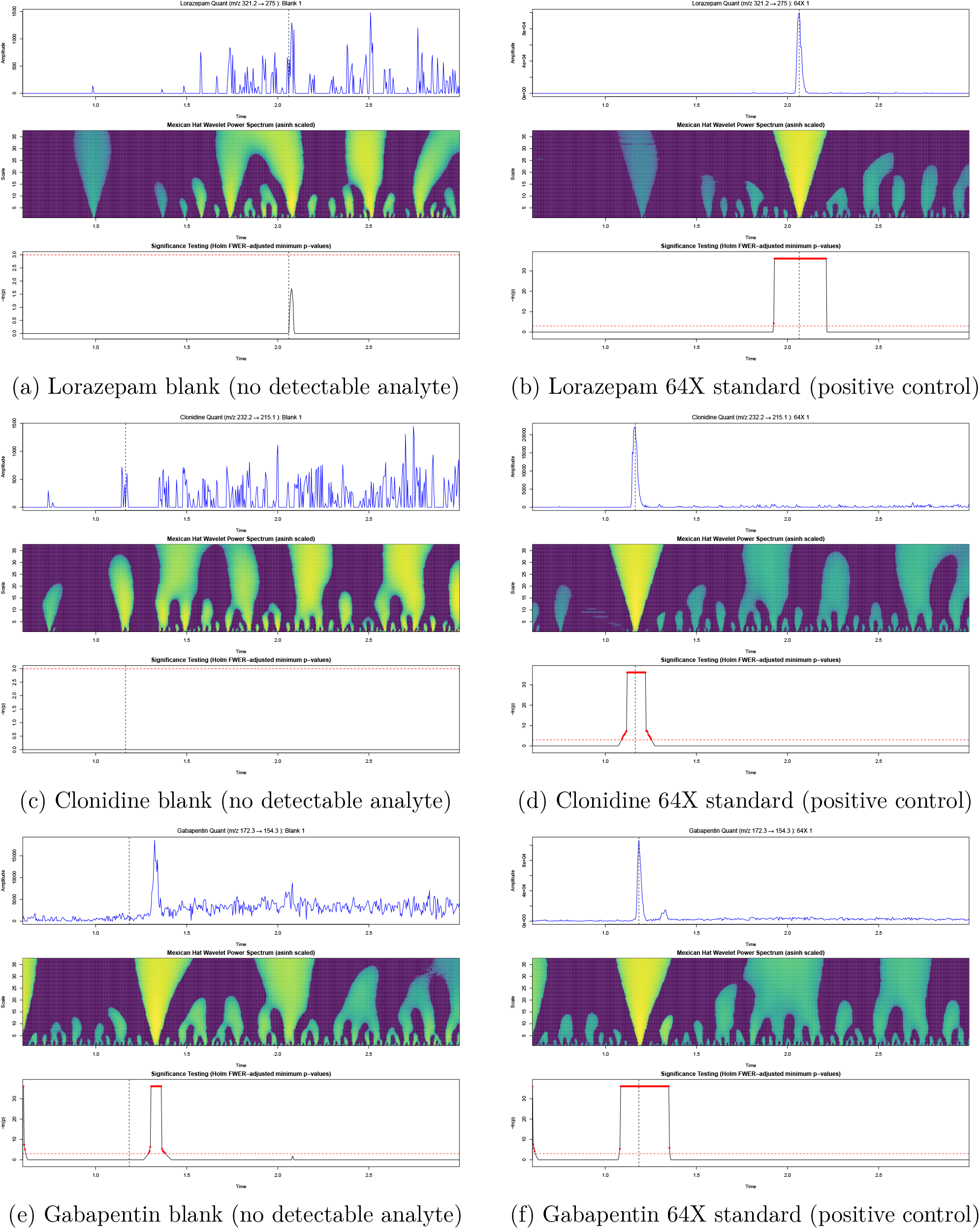
Chromatograms and wavelet-based significance maps for the quantifier transitions of Lorazepam, Clonidine, and Gabapentin on the Waters Xevo TQ Absolute XR. Left column: blank plasma. Right column: 64X standards (5.00 ng/mL).

Figure 4a-f shows the quantifier SRM transitions (Table 1) for all three compounds in the dilution-series validation. In the 64X standards, each compound shows a strong, symmetric peak at the internal-standard-aligned retention time, with DOG2 wavelet power forming a sharp multiscale ridge and a contiguous region of Holm-adjusted *p*_adj_ ≪ 0.05 (Figure 4b, d, f). By contrast, the blank plasma samples (Figure 4a, c, e) show only low-level baseline structure that our Monte Carlo test classified as compatible with the chemical-noise null, confirming the specificity of the wavelet-based detection.

Table 2 summarises the detection results across the full dilution series for all three compounds. Detection is defined as at least one significant time point (*p*_adj_ *<* 0.05) within ±0.1 min of the internal-standard-aligned retention time (Algorithm 2). All three compounds produced zero false positives in blanks. Lorazepam, Clonidine, and Gabapentin had empirical LODs of 2X (0.156 ng/mL), 8X (0.625 ng/mL), and 4X (0.313 ng/mL), respectively, defined as the lowest concentration at which all three replicates were detected. Note that a baseline-separated chemical-noise peak near the Gabapentin retention time is present in blank and low-concentration samples but is resolved from the analyte peak and was not counted as a detection.

The three compounds span a range of detection sensitivity, with empirical LODs from 2X (Lorazepam) to 8X (Clonidine), reflecting differences in ionization efficiency, transition selectivity, and matrix effects across the three SRM channels. At concentrations near the empirical LOD, partial detection (1/3 or 2/3 replicates) is observed for Lorazepam (0.5X-1X) and Gabapentin (2X). This transition-zone behavior is expected: as the analyte signal approaches the noise floor, statistical power declines and detection becomes stochastic rather than deterministic. Above the LOD, detection is consistent (3/3) for all compounds and concentrations.

As further external validation, the wavelet method was applied to an independent clinical pain panel that used a SCIEX 6500 to measure ketamine in urine with a LLOQ of 2ng/mL (m/z 238→ 125; 16 samples, same 10^6^ simulations, and 5% FWER). The low calibration standard showed 10 significant time points; the matrix blank showed no significant features; and four biological samples with confirmed sub-LLOQ ketamine showed 4-10 significant time points at the expected retention time. The remaining nine biological negatives and one sample containing chemical noise at an incorrect retention time were correctly classified as non-significant. The full results and example data are provided in Table S4 and Figures S1-S6.

### Example: Analysis of Novel SPM Identification

To be useful in difficult detection applications, the wavelet method has to work across instrument makes and models as well as compound classes. The dilution-series validation was performed on data from a Waters instrument measuring drugs in plasma with known concentrations. A more challenging application is the detection of specialized pro-resolving mediators (SPMs). These lipids are often reported at pg/mL concentrations^28^. Detection of SPMs by targeted SRM LC-MS/MS is currently under active discussion in the literature, with particular attention being focused on reporting standards, the role of S/N criteria, and whether low-level signals in SRM channels constitute sufficient evidence for identification^2931^. Walker et al. (2024)^15^ reported the detection of resolvin T4 (RvT4) in plasma and deposited the raw instrument files in a public repository, providing a direct link between an identification claim and the underlying SRM data. Because the measurements were made in plasma and the raw data are publicly available, the Walker et al. dataset is a suitable test case for our wavelet-based approach.

We obtained the raw LC-MS/MS data via repository accession S-BSST880^32^. The published dataset comprises raw data files containing data from samples analyzed for RvT4 and related specialized pro-resolving mediators (SPMs).

Plasma RvT4 samples were analyzed in two separate experiments. The file MW-RvT4ex3-plasma MS4-MF was collected using a Sciex 6500 and contains eight plasma samples (four Chow, four Western diet). The file JD ACP ApoE RvT 052022 was collected using a Sciex 7500 and contains six plasma samples (three Chow, three Western diet). Sample names and group assignments for all acquisitions follow the information in Walker et al. (2024) Supplemental table (Sample legend.docx). Table S3 summarizes the raw data files, instruments, and SRM transitions analyzed

Vendor raw files were converted to the open mzML format using MSConvert (ProteoWizard)^33^ version 3.0.24018. SCIEX OS software version 3.3 (SCIEX, Framingham, MA, USA) was used for checking instrument vendor files and sample metadata. The raw file names (wiff, wiff.scan, wiff2) were listed in the Walker et al. (2024) supplemental information. All subsequent wavelet, Monte Carlo, and detection analyses were performed on the converted mzML files.

For both Sciex instruments, the RvT4 quant blank intensities showed no evidence of left-censoring, and Shapiro-Wilk tests were consistent with normality (6500: *p* = 0.065; 7500: *p* = 0.16), so ordinary sample mean and standard deviation were used directly for the electronic-noise parameters rather than the ROS procedure required for the Waters data.

### Quantifier transitions for RvT4

We applied the wavelet framework to the RvT4 quantifier transition (361.1→211.1; Table S3) across all 14 plasma samples (8 from the 6500 run, 6 from the 7500 run; see Table S2). Figure 5a-b shows RvT4 standards on each instrument. In both cases, the standards exhibit tall, well-defined peaks at the expected retention time with intense multiscale wavelet ridges and extremely small adjusted *p*-values, confirming that the assays can readily detect the RvT4 molecule.

**Figure 5.**
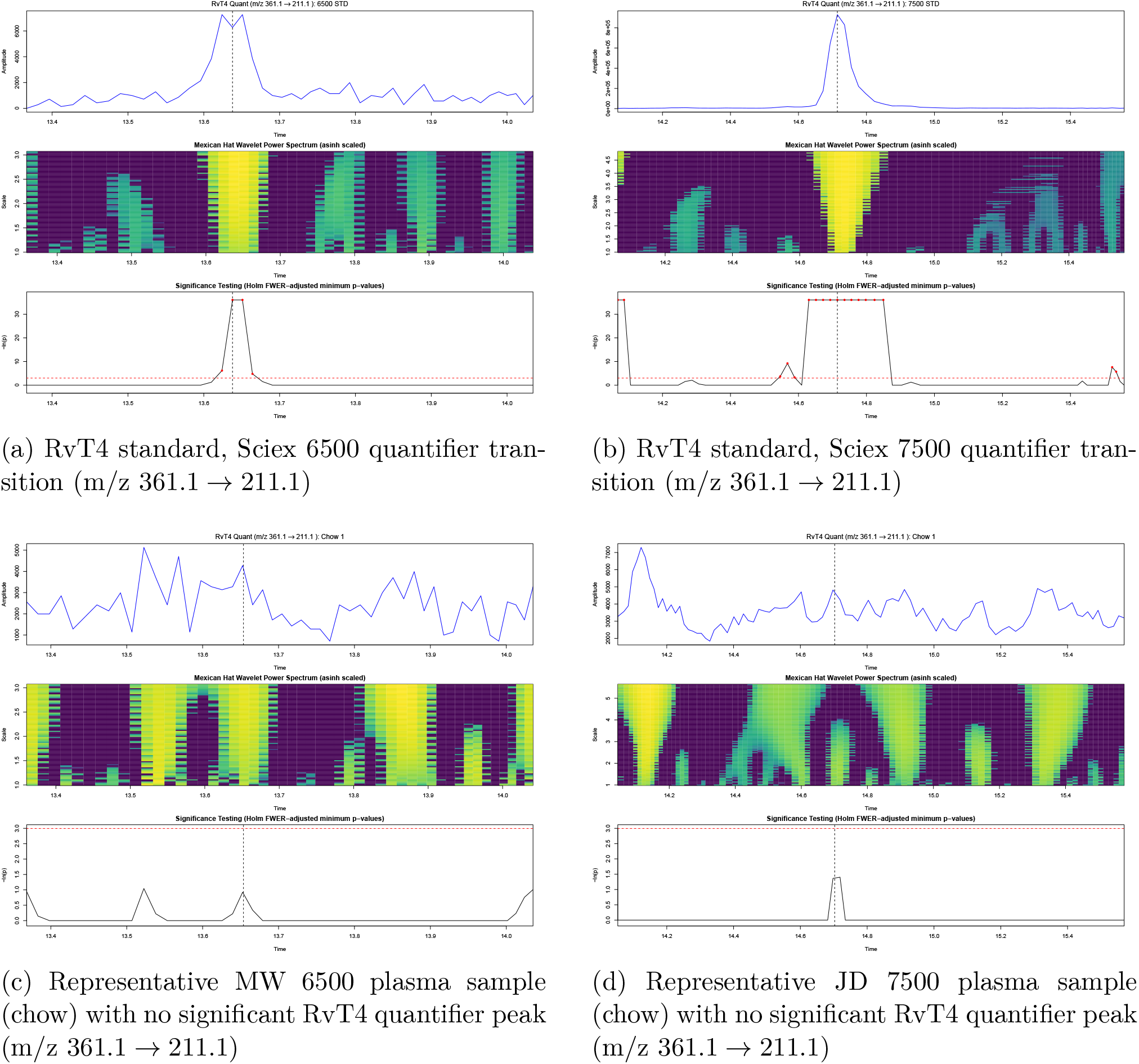
RvT4 quantifier ion (m/z 361.1→ 211.1) on two instruments. (a-b) Calibration standards on the Sciex 6500 and 7500. (c-d) Representative Chow diet plasma samples from the MW 6500 and JD 7500 runs. Variations in the time axis reflect different LC-MS/MS methods (Table S1).

The behaviour in biological plasma samples is markedly different (Figure 5c-d). Across the 8 MW (6500) RvT4 chromatograms, most traces resemble the blanks: low-level, irregular structure with several modest peaks scattered across the chromatogram. In the wavelet domain, this produces isolated islands of power, but the minimum-*p* traces remain below the significance threshold at the RvT4 retention time in all 8 of the samples. On the 7500 JD run (6 plasma samples), none of the chromatograms display a significant RvT4 quantifier peak at the expected retention time after Holm-Bonferroni correction.

Of the 14 plasma samples analyzed, Walker et al. reported quantified RvT4 concentrations in 7 (5 Chow, 2 Western diet; Table S2). Our Monte Carlo procedure shows that none of the 14 samples, including the 7 originally reported as RvT4-positive, contains a statistically significant quantifier peak at the expected retention time. Per-sample chromatograms, wavelet power maps, and significance test results for all standards and plasma samples from both instruments are available in the project repository (see Supporting Information).

## Discussion

### Principal Findings

We present a wavelet-based Monte Carlo method for objective SRM peak detection in the presence of structured chemical noise and chemical interference. The method operates at the level of individual time points and wavelet scales, assigning per-time-point *p*-values by comparing multiscale wavelet power to an empirical null model. Holm-Bonferroni correction across time points controls the FWER at a user-specified level for each chromatogram.

In a three-compound dilution series with known ground truth (Lorazepam, Clonidine, Gabapentin in K_2_EDTA plasma), the method produced zero false-positive detections in blanks across all compounds while achieving consistent detection (3/3 replicates) at and above each compounds empirical LOD. Partial detection in transition-zone concentrations just below the LOD is consistent with declining statistical power near the noise floor. The validation spanned two vendor platforms (Waters Xevo TQ Absolute XR for the dilution series; Sciex 6500 and 7500 for the SPM data), suggesting portability across instruments and compound classes.

In a published SRM LC-MS/MS study targeting RvT4, the Monte Carlo wavelet approach robustly detects standards while all low-level features in the RvT4 channels are consistent with the chemical-noise background when evaluated using an adjusted *p*-value threshold of 0.05.

### Limitations and Future Directions

Our implementation has several limitations that should be considered when interpreting the current results and when applying the method to other datasets.

Null-model specification and use of blanks: The null model has two distinct components, chemical noise and electronic noise, with different data requirements. The chemical-noise component is built from peaks detected outside the analyte exclusion window across all available chromatograms, including study samples. Matrix blanks are not required: the exclusion window around the expected analyte retention time prevents true signal from contaminating the noise distributions, so the chemical-noise model can be constructed entirely from study samples. When matrix blanks are available, their chemical-noise peaks are simply pooled with those from the study samples, improving the sampling of the noise distributions.

The electronic-noise component, by contrast, does require a blank, but only a solvent or reagent blank, not a matrix-matched blank. Electronic noise is a characteristic of the instrument and detector, not of the sample matrix. In the present work we estimated the electronic-noise parameters (*µ*_blank_, *σ*_blank_) from reagent-blank injections using regression on order statistics to account for left-censoring at zero intensity. Any method that provides comparable estimates of the electronic noise would suffice, including using normal statistics on reagent blanks if they meet acceptable criteria of normality.

If matrix blanks are available and used, two caveats apply. If blanks are systematically cleaner than study samples, or if sample-specific interferences occur preferentially in the analyte window (for example, because of co-eluting matrix components), then the null model may underestimate the true false-positive rate. Conversely, extremely noisy blanks could make the test overly conservative. Future work should examine how sensitive the Monte Carlo thresholds are to the choice and number of samples used to characterize the chemical noise distributions. Hierarchical models that share information across instruments and studies, or that explicitly allow for rare high-impact interferences in specific retention-time regions, should also be considered.

Stability of chemical noise: Relatedly, we assume that the pattern of chemical noise for a given instrument and transition is roughly stable over the time span of each acquisition series. This may break down in long-running studies or when substantial changes in chromatography, source tuning, or sample preparation occur. Extending the model to allow slowly varying background structure, e.g., via time-varying mixtures of empirical peak noise profiles or by stratifying the null by batch, would make the framework more robust to such drift.

Computational cost and scalability: The current implementation performs up to 10^6^ Monte Carlo simulations per chromatogram. While this cost is acceptable for the validation and retrospective analyses presented here, the obvious step is to move away from R and use highly parallel Monte Carlo implementations (GPU or cluster) since each simulation is independent.

Scope of application: We validated the method exclusively on SRM chromatograms of drugs spiked into plasma and applied it to targeted SRM lipidomics data. We did not attempt to combine evidence across multiple transitions or samples in a formal hierarchical testing framework. Incorporating joint models for related transitions, or combining power across related transitions, may reduce false positives while preserving rigorous error control.

Looking ahead, we envision several extensions. Packaging the workflow into an open-source library could simplify adoption, although the full source code and analysis inputs are already provided for reproducibility. Prospective application to new SPM and lipid mediator studies alongside orthogonal confirmation methods would help define practical decision rules for declaring the presence or absence of low-abundance analytes. Because the method operates on any one-dimensional chromatographic trace in which a specific analyte is claimed at a defined retention time, it could in principle be applied to extracted ion chromatograms from high-resolution instruments, though this remains to be validated. Full-scan or data-independent acquisition modes will require additional work to handle non-stationary chemical noise and more complex background structure. Finally, adapting the approach to other analytical contexts such as metabolomics, environmental trace analysis, or MALDI imaging may provide a general path toward CFAR-like detection wherever chemical noise dominates traditional electronic noise.

## Conclusion

There are cases where trace-level SRM LC-MS/MS measurements are more limited by structured chemical noise rather than by electronic fluctuations. By modeling chemical noise empirically and using wavelet-based Monte Carlo significance testing with family-wise error control, we turn visual peak-calling heuristics into explicit hypothesis tests. In a three-compound dilution series with known ground truth, the method correctly detected analytes at and above their empirical LODs with zero blank false positives, and exhibited the expected gradual loss of power in the transition zone near each LOD. In an independent ketamine pain panel, the method detected four confirmed positive samples with signal intensities below the low calibration standard. In the Walker et al. (2024) RvT4 dataset, this framework aligns with chemical intuition for positive controls while indicating that, under our empirical-null model, putative detections of a novel mediator are statistically indistinguishable from interference. More broadly, the results imply that credible claims about low-abundance analytes require robust noise characterization and additional confirmatory transitions to substantiate claims of detection in the presence of structured chemical noise. We expect that adopting similar CFAR-like frameworks in SRM LC-MS/MS will materially reduce false positives in trace analysis and sharpen the evidentiary standards for detection claims.

## Supporting Information

Supplementary data contains the RvT4 reanalysis data tables from Walker et al. (2024) and the ketamine external validation results (Table S4, Figures S1-S6). All source code, mzML data files, and per-sample analysis results for both the dilution-series and RvT4 datasets are available at https://github.com/rkjulian/wavelet-peak-significance.

## Acknowledgments

Thanks to Dr. Oscar Waddell for thoughtfully reading and providing comments on early drafts of this manuscript. The authors used large language models (ChatGPT, Claude, and others) for assistance with text editing, plotting syntax assistance (axis alignment, legend placement, panel arrangement), and LaTeX formatting help. All scientific content, data analysis, and figures were created by the authors.

## Author Contributions

R.K.J. developed the wavelet Monte Carlo algorithm and code. B.A.R. analyzed the published SPM data. S.R.M. and F.Y. designed and performed the LC-MS/MS drug dilution experiment. All authors contributed equally to the study design, data analysis, and manuscript writing.

## For Table of Contents Only

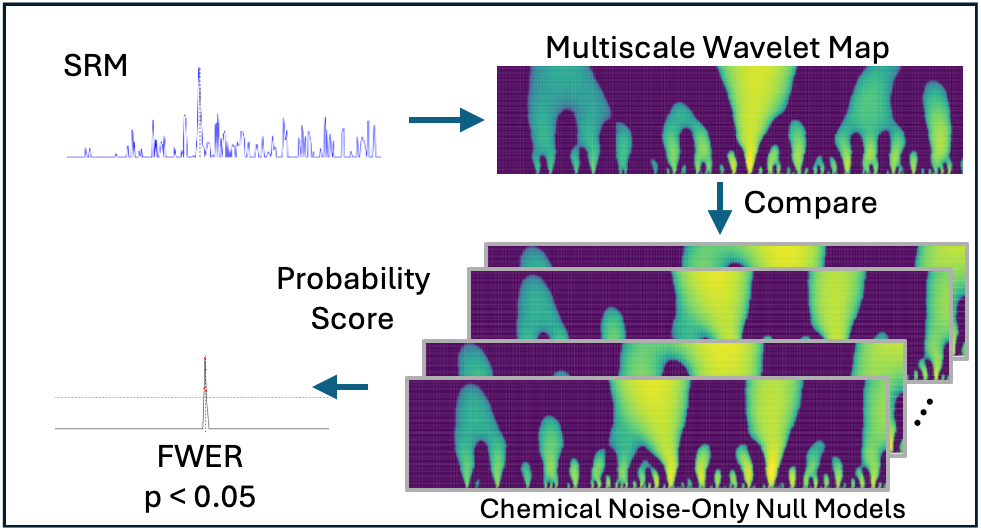

## References

(1) Bonfiglio, R.; King, R. C.; Olah, T. V.; Merkle, K. The Effects of Sample Preparation Methods on the Variability of the Electrospray Ionization Response for Model Drug Compounds. Rapid Communications in Mass Spectrometry 1999, 13 (12), 11751185. 10.1002/(SICI)1097-0231(19990630)13:12%3C1175::AID-RCM639%3E3.0.CO;2-0.

(2) Kushnir, M. M.; Rockwood, A. L.; Nelson, G. J.; Yue, B.; Urry, F. M. Assessing Analytical Specificity in Quantitative Analysis Using Tandem Mass Spectrometry. Clinical Biochemistry 2005, 38 (4), 319327. 10.1016/j.clinbiochem.2004.12.003.

(3) Guo, X.; Bruins, A. P.; Covey, T. R. Characterization of Typical Chemical Background Interferences in Atmospheric Pressure Ionization Liquid Chromatography-Mass Spectrometry. Rapid Communications in Mass Spectrometry 2006, 20 (20), 31453150. 10.1002/rcm.2715.

(4) Trötzmüller, M.; Guo, X.; Fauland, A.; Köfeler, H.; Lankmayr, E. Characteristics and Origins of Common Chemical Noise Ions in Negative ESI LCMS. Journal of Mass Spectrometry 2011, 46 (6), 553560. 10.1002/jms.1924.

(5) Guo, X.; Bruins, A. P.; Covey, T. R.; Trötzmüller, M.; Lankmayr, E. Alternative Reagents for Chemical Noise Reduction in Liquid Chromatography-Mass Spectrometry Using Selective Ion-Molecule Reactions. Journal of the American Society for Mass Spectrometry 2009, 20 (1), 105111. 10.1016/j.jasms.2008.09.021.

(6) Krutchinsky, A. N.; Chait, B. T. On the Nature of the Chemical Noise in MALDI Mass Spectra. Journal of the American Society for Mass Spectrometry 2002, 13 (2), 129134. 10.1016/S1044-0305(01)00336-1.

(7) Kumler, W.; Hazelton, B. J.; Ingalls, A. E. Picky with Peakpicking: Assessing Chromatographic Peak Quality with Simple Metrics in Metabolomics. BMC Bioinformatics 2023, 24 (1), 404. 10.1186/s12859-023-05533-4.

(8) Vangeenderhuysen, P.; Vynck, M.; Pomian, B.; De Windt, K.; Callemeyn, E.; De Paepe, E.; De Commer, L.; Raes, J.; Nawrot, T.; Rainer, J.; Hemeryck, L. Y.; Vanhaecke, L. Automated Integration and Quality Assessment of Chromatographic Peaks in LCMS-Based Metabolomics and Lipidomics Using TARDIS. Analytical Chemistry 2025, 97 (18), 99279934. 10.1021/acs.analchem.5c00567.

(9) Hope, A. C. A. A Simplified Monte Carlo Significance Test Procedure. Journal of the Royal Statistical Society: Series B (Methodological) 1968, 30 (3), 582598. 10.1111/j.2517-6161.1968.tb00759.x.

(10) Holm, S. A Simple Sequentially Rejective Multiple Test Procedure. Scandinavian Journal of Statistics 1979, 6 (2), 6570.

(11) Cappadona, S.; Levander, F.; Jansson, M.; James, P.; Cerutti, S.; Pattini, L. Wavelet-Based Method for Noise Characterization and Rejection in High-Performance Liquid Chromatography Coupled to Mass Spectrometry. Analytical Chemistry 2008, 80 (13), 49604968. 10.1021/ac800166w.

(12) Dang, M. C.; Patil, A. A.; Li, T. K. L.; Chou, S.-W.; Thi Hoang, T. K.; Estayan, M. I. C.; Peng, W.-P. Optimization of the Entropy-Based Wavelet Method for Removing Strong RF and AC Interferences in a Charge Detection Linear Ion Trap Mass Spectrometer. Analytical Chemistry 2025, 97 (9), 50665076. 10.1021/acs.analchem.4c06069.

(13) Maraun, D.; Kurths, J. Cross Wavelet Analysis: Significance Testing and Pitfalls. Nonlinear Processes in Geophysics 2004, 11 (4), 505514. 10.5194/npg-11-505-2004.

(14) Rohling, H. Radar CFAR Thresholding in Clutter and Multiple Target Situations. IEEE Transactions on Aerospace and Electronic Systems 1983, AES-19 (4), 608621. 10.1109/TAES.1983.309350.

(15) Walker, M. E.; De Matteis, R.; Perretti, M.; Dalli, J. Resolvin T4 Enhances Macrophage Cholesterol Efflux to Reduce Vascular Disease. Nature Communications 2024, 15 (1), 975. 10.1038/s41467-024-44868-1.

(16) R Core Team. R: A Language and Environment for Statistical Computing; R Foundation for Statistical Computing: Vienna, Austria, 2025.

(17) Gatto, L.; Rainer, J.; Gibb, S. MSnbase: Base Functions and Classes for Mass Spectrometry and Proteomics; 2025. 10.18129/B9.bioc.MSnbase.

(18) Gatto, L.; Lilley, K. MSnbase - an r/Bioconductor Package for Isobaric Tagged Mass Spectrometry Data Visualization, Processing and Quantitation. Bioinformatics 2012, 28, 288289.

(19) Gatto, L.; Gibb, S.; Rainer, J. MSnbase, Efficient and Elegant r-Based Processing and Visualisation of Raw Mass Spectrometry Data. bioRxiv 2020.

(20) Du, P.; Kibbe, W.; Lin, S.; Oller Moreno, S. MassSpecWavelet: Peak Detection for Mass Spectrometry Data Using Wavelet-Based Algorithms; 2025. 10.18129/B9.bioc.MassSpecWavelet.

(21) Du, P.; Kibbe, W. A.; Lin, S. M. Improved Peak Detection in Mass Spectrum by Incorporating Continuous Wavelet Transform-Based Pattern Matching. Bioinformatics 2006, 22 (17), 20592065. 10.1093/bioinformatics/btl355.

(22) Lee, L. NADA: Nondetects and Data Analysis for Environmental Data; 2025. 10.32614/CRAN.package.NADA.

(23) Mersmann, O.; Trautmann, H.; Steuer, D.; Bornkamp, B. Truncnorm: Truncated Normal Distribution; 2023.

(24) Torrence, C.; Compo, G. P. A Practical Guide to Wavelet Analysis. 1998.

(25) Ripley, B. D. Stochastic Simulation; Wiley series in probability and mathematical statistics. Applied probability and statistics; Wiley: New York, 1987. 10.1002/9780470316726.

(26) Helsel, D. R. Nondetects and Data Analysis: Statistics for Censored Environmental Data; Statistics in practice; Wiley-Interscience: Hoboken, N.J., 2005.

(27) Mallat, S.; Hwang, W. L. Singularity Detection and Processing with Wavelets. IEEE Transactions on Information Theory 1992, 38 (2), 617643. 10.1109/18.119727.

(28) Kutzner, L.; Rund, K. M.; Ostermann, A. I.; Hartung, N. M.; Galano, J.-M.; Balas, L.; Durand, T.; Balzer, M. S.; David, S.; Schebb, N. H. Development of an Optimized LC-MS Method for the Detection of Specialized Pro-Resolving Mediators in Biological Samples. Frontiers in Pharmacology 2019, 10, 169. 10.3389/fphar.2019.00169.

(29) ODonnell, V. B.; FitzGerald, G. A.; Murphy, R. C.; Liebisch, G.; Dennis, E. A.; Quehenberger, O.; Subramaniam, S.; Wakelam, M. J. O. Steps Toward Minimal Reporting Standards for Lipidomics Mass Spectrometry in Biomedical Research Publications. Circulation. Genomic and Precision Medicine 2020, 13 (6), e003019. 10.1161/CIRCGEN.120.003019.

(30) Dalli, J.; Gomez, E. A.; Serhan, C. N. Evidence for the Presence and Diagnostic Utility of SPM in Human Peripheral Blood, 2022. 10.1101/2022.04.28.489064.

(31) ODonnell, V. B.; Schebb, N. H.; Milne, G. L.; Murphy, M. P.; Thomas, C. P.; Steinhilber, D.; Gelhaus, S. L.; Kühn, H.; Gelb, M. H.; Jakobsson, P.-J.; Blair, I. A.; Murphy, R. C.; Freeman, B. A.; Brash, A. R.; FitzGerald, G. A. Failure to Apply Standard Limit-of-Detection or Limit-of-Quantitation Criteria to Specialized Pro-Resolving Mediator Analysis Incorrectly Characterizes Their Presence in Biological Samples. Nature Communications 2023, 14 (1), 7172. 10.1038/s41467-023-41766-w.

(32) BioStudies. BioStudies < The European Bioinformatics Institute < EMBL-EBI. https://www.ebi.ac.uk/biostudies/studies/S-BSST880?query=S-BSST880 (accessed 2024-04-06).

(33) Kessner, D.; Chambers, M.; Burke, R.; Agus, D.; Mallick, P. ProteoWizard: Open Source Software for Rapid Proteomics Tools Development. Bioinformatics 2008, 24 (21), 25342536. 10.1093/bioinformatics/btn323.

