## Supplemental Information for "Monte Carlo Wavelet Analysis for Objective Peak Detection in SRM LC-MS/MS Analysis"

May 20, 2026

[1]Indigo BioAutomation, Inc., Carmel, IN 46032, USA
[2]Department of Chemistry, Purdue University, West Lafayette, IN 47907, USA
[3]Laboratory Corporation of America Holdings, Research Triangle Park, NC 27709, USA
[4]School of Health Sciences, University of Iceland, 102 Reykjavik, Iceland
[5]Center for Diagnostic Innovation (CDI), Children's Hospital of Philadelphia, Philadelphia, PA 19104, USA
[6]Department of Pathology and Laboratory Medicine, Children's Hospital of Philadelphia, Philadelphia, PA 19104, USA
[7]Perelman School of Medicine, University of Pennsylvania, Philadelphia, PA 19104, USA

#### Contents

This Supplemental Information document provides details of the RvT4 lipid mediator re-analysis and the ketamine external validation used as demonstrations in the primary article.

- **Table S1.** Methodological variations across the reanalyzed RvT4 dataset from Walker et al. (2024)
- **Table S2.** Number of quantified RvT4 peaks by diet group as reported in Walker et al. (2024)
- **Table S3.** Summary of LC-MS/MS data files and transitions reanalyzed from Walker et al. (2024)
- **Table S4.** Ketamine external validation: wavelet significance test results
- **Figure S1.** Ketamine low standard (STD) significance test
- **Figure S2.** Ketamine blank significance test
- **Figure S3.** Ketamine sub-LLOQ biological positive (Inj 27) significance test
- **Figure S4.** Ketamine biological negative (Inj 19) significance test
- **Figure S5.** Ketamine chemical noise characterization: STD
- **Figure S6.** Ketamine chemical noise characterization: Inj 19

### RvT4 Reanalysis: Data and Methods

Different LC and MS methods were employed across the various runs published in Walker et al. (2024). These included changes in the SRM acquisition settings, multiple instruments (Sciex 6500 and 7500), variations in flow rate (e.g., 350 uL/min for JD ACP ApoE RvT 052022), and the LC platform. These methodological variations are detailed in Table S1 and account for observed differences in retention times and peak characteristics between samples.

Table S1. Methodological variations across the reanalyzed dataset from Walker et al. (2024). These differences account for variations in retention times and peak characteristics observed in Table S3 and in the main text. All samples were analyzed in negative ionization mode with selected reaction monitoring (SRM).

| File name | Run date | LC system | MS method file |
|---|---|---|---|
| Mary RvT4 in IA aorta MS4 | 2018-11-28 | Shimadzu | All dIS-Full Neg Indiv-sMRM.dam |
| JD ACP MW ART 051822 | 2018-05-18 | Shimadzu | All dIS-Full Neg Indiv-sMRM long window-1120.dam |
| MWRvT4ex3-plasma MS4-MF | 2019-05-30 | Shimadzu | All dIS-Full Neg Indiv-sMRM.dam |
| JD ACP ApoE RvT 052022 | 2022-05-20 | Exion | Negative mode Profiling.dam - modified flow 052022.dam |

In Walker et al. (2024) quantified values are given for plasma RvT4 samples measured across two experiments from 14 samples. The number of reported detections by diet group is shown in Table S2.

Table S2. Number of quantified RvT4 peaks (from 14 total samples) by diet as reported in Walker et al. (2024).

| Diet | RvT4 |
|---|---|
| Chow | 5 |
| WD | 2 |

Table S3. Summary of LC-MS/MS data files and transitions reanalysed from Walker et al. (2024). Points is the number of acquired data points (time measurements) in each chromatogram; this value determines the maximum wavelet scale via $s_{\max} = \min\left((N-1)/16,\, 64\right)$ (see Methods).

| Sample Type | Compound | File name | Instrument | Chrom. | Transition | Points |
|---|---|---|---|---|---|---|
| Plasma | RvT4 | MW-RvT4ex3-plasma MS4-MF | Sciex 6500 | Quant | 361.1 → 211.1 | 50 |
| | | | | IS | 339.2 → 197.2 | 51 |
| Plasma | RvT4 | JD ACP ApoE RvT 052022 | Sciex 7500 | Quant | 361.1 → 211.1 | 91 |
| | | | | IS | 339.2 → 197.2 | 91 |

Per-sample chromatograms, wavelet power maps, chemical noise characterizations, and significance test results for all RvT4 compounds and transitions are available in the project repository at https://github.com/rkjulian/wavelet-peak-significance.

### Ketamine External Validation

As additional external validation, the Monte Carlo wavelet significance method was applied to an independent ketamine clinical pain panel assay acquired on a SCIEX 6500 instrument. This dataset is entirely separate from both the Waters dilution-series validation and the RvT4 reanalysis, representing a third compound class (anesthetic/analgesic) on the same SCIEX platform family. The ketamine quantifier transition (m/z 238 $\rightarrow$ 125) was analyzed across 16 samples: one low calibration standard (STD), one matrix blank, and 14 biological samples. The internal standard eluted at approximately 2.13 min, with the ketamine quantifier peak at approximately 1.94 min (offset $-0.193$ min). All analyses used $10^6$ Monte Carlo simulations with Holm–Bonferroni FWER correction at $\alpha = 0.05$.

Table S4 summarizes the wavelet significance test results for all 16 ketamine samples.

Table S4. Ketamine external validation: wavelet significance test results. The quantifier transition (m/z 238 $\rightarrow$ 125) was analyzed on a SCIEX 6500 with $10^6$ Monte Carlo simulations at $\alpha = 0.05$.

| Sample | Sig. Time Points | Min $p_{\text{adj}}$ [1] | Result |
|---|---|---|---|
| STD | 10 | $< 2.22 \times 10^{-16}$ | Detected (low standard) |
| Blank | 0 | 0.613 | Not detected (true negative) |
| Inj 7 | 0 | 1.000 | Not detected |
| Inj 8 | 7 | $< 2.22 \times 10^{-16}$ | Detected (sub-LLOQ positive) |
| Inj 11 | 7 | $< 2.22 \times 10^{-16}$ | Detected (sub-LLOQ positive) |
| Inj 12 | 0 | 0.694 | Not detected |
| Inj 13 | 10 | $< 2.22 \times 10^{-16}$ | Detected (sub-LLOQ positive) |
| Inj 14 | 0 | 0.405 | Not detected |
| Inj 15 | 0 | 1.000 | Not detected |
| Inj 16 | 0 | 0.685 | Not detected |
| Inj 19 | 0 | 0.993 | Not detected |
| Inj 21 | 0 | 0.886 | Not detected |
| Inj 22 | 0 | 0.978 | Not detected |
| Inj 24 | 0 | 0.989 | Not detected |
| Inj 25 | 0 | 0.149 | Not detected (chemical noise) |
| Inj 27 | 4 | $1.01 \times 10^{-2}$ | Detected (sub-LLOQ positive) |

[1] $2.22 \times 10^{-16}$ is the smallest representable difference between two double-precision floating-point numbers in R (`.Machine$double.eps`).

The low calibration standard (STD) exhibits a well-defined peak at the expected retention time with 10 contiguous significant time points ($p_{\text{adj}} < 2.22 \times 10^{-16}$; Figure S1), confirming that the assay reliably detects ketamine even at the lowest calibrator level. The blank shows no significant features, with a minimum adjusted $p$-value of 0.613 (Figure S2), confirming specificity.

Among the 14 biological samples, four (Inj 8, 11, 13, 27) are confirmed ketamine-positive with signal intensity below the low calibration standard. The wavelet significance test detects all four, with 4–10 significant time points centered at the expected retention time. Inj 27 (Figure

S3) is a representative sub-LLOQ positive with 4 significant time points ($p_{\text{adj}} = 1.01 \times 10^{-2}$), demonstrating that the method detects analyte signal below conventional reporting limits. The remaining nine biological negatives show no significant detections.

Inj 25 is an instructive case. Its chromatogram contains real signal, but at a retention time inconsistent with the expected ketamine peak. The Monte Carlo test correctly classifies this as non-significant ($p_{\text{adj}} = 0.149$), consistent with chemical noise rather than analyte.

Inj 19 (Figure S4) is representative of the biological negative samples whose chromatographic features are visually similar to the types of peaks that might be manually integrated in conventional analysis. The chemical noise characterization (Figure S6) shows the wavelet power spectrum for this sample, illustrating the level of structured chemical background against which the significance test operates. This contrasts with the STD chemical noise profile (Figure S5), where the dominant feature aligns precisely with the expected retention time.

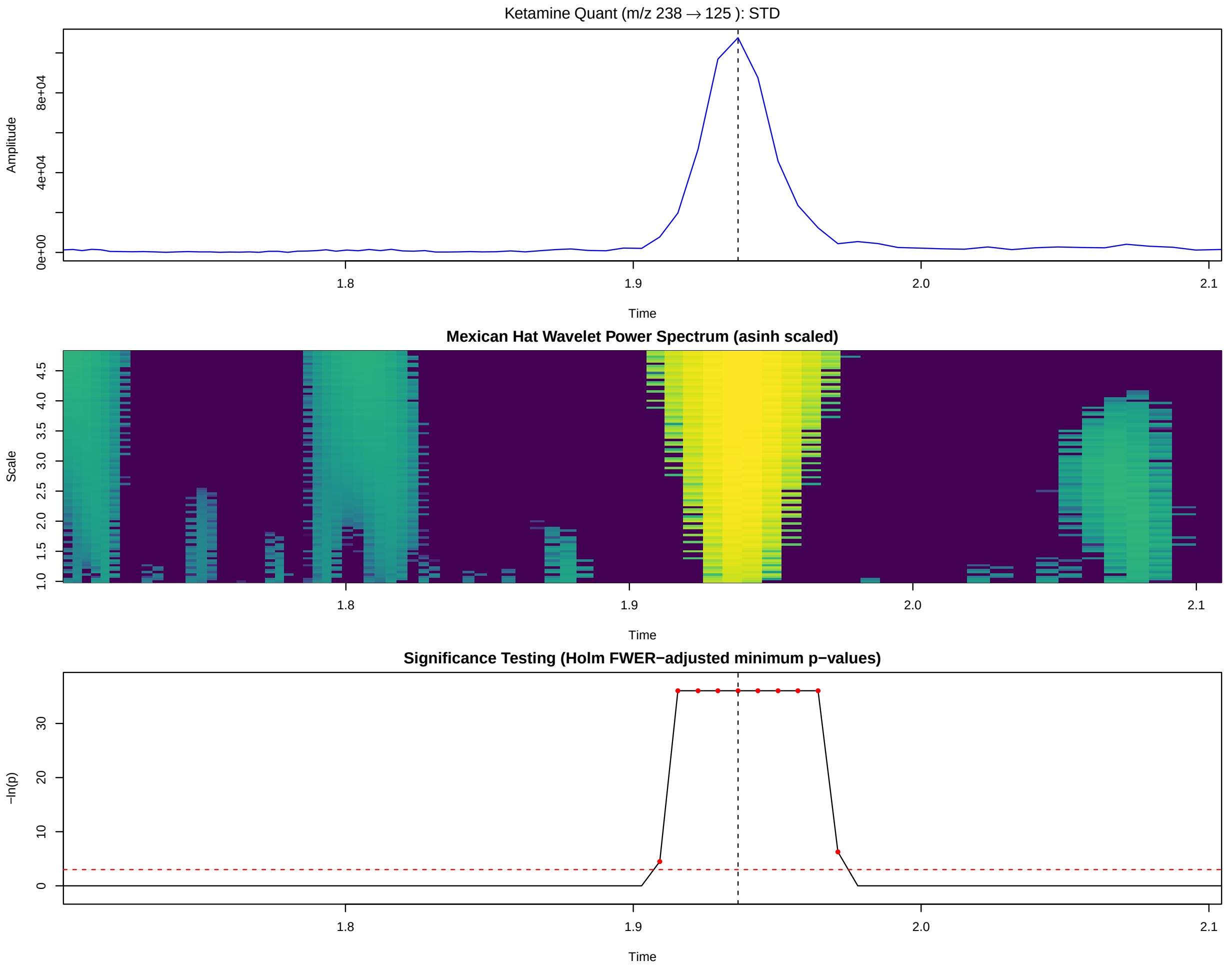


Figure S1. Ketamine low standard (STD) significance test on the SCIEX 6500. Quantifier transition m/z 238 → 125. Top: chromatogram with expected retention time (dashed line). Middle: Mexican Hat wavelet power spectrum (asinh scaled). Bottom: Holm–Bonferroni adjusted $p$-values with significance threshold at $-\ln(0.05)$ (red dashed line). The low calibration standard shows 10 contiguous significant time points ($p_{\mathrm{adj}} < 2.22 \times 10^{-16}$) at the expected retention time.

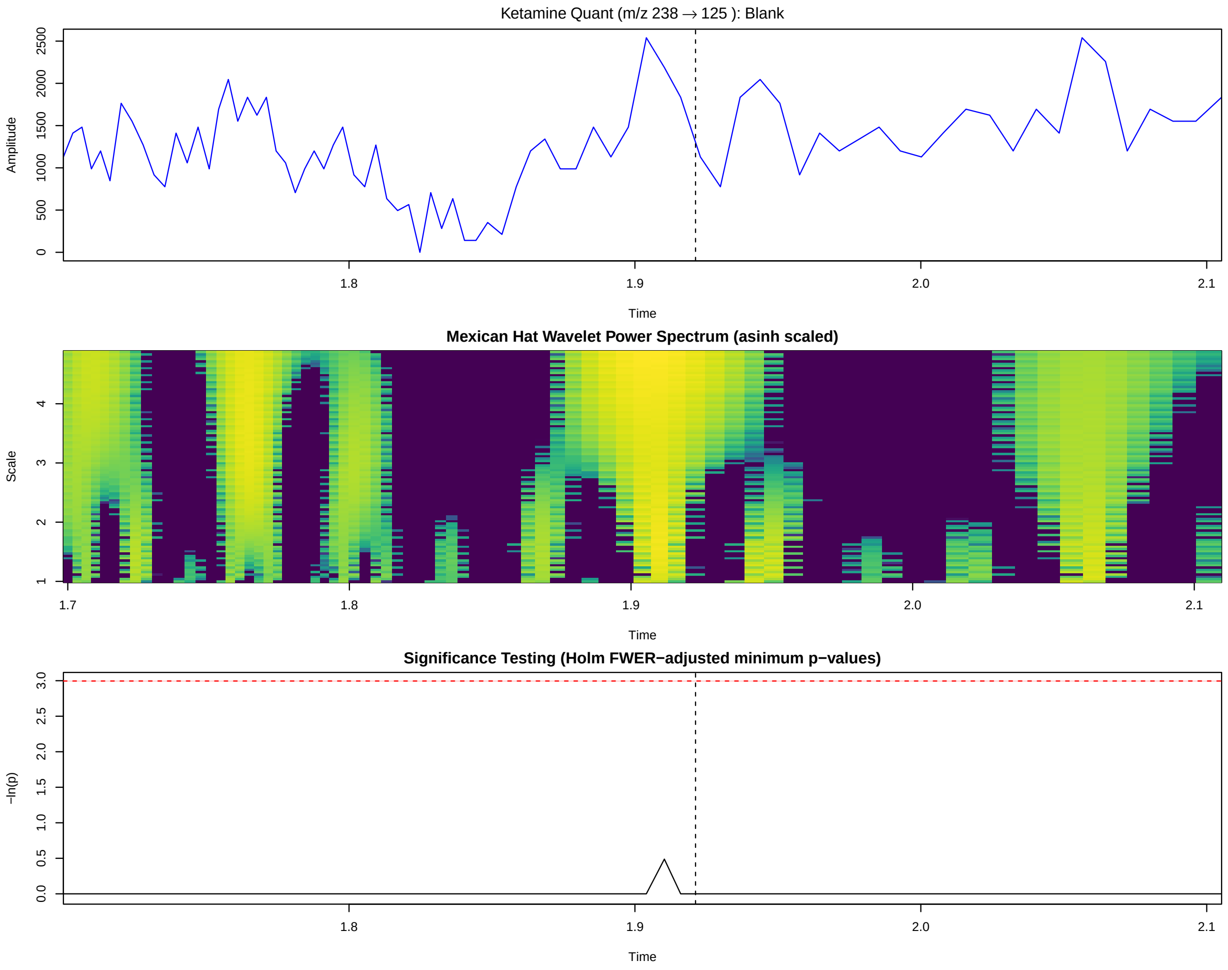


Figure S2. Ketamine blank significance test on the SCIEX 6500. Quantifier transition m/z 238 → 125. No significant features detected; minimum adjusted $p$-value = 0.613.

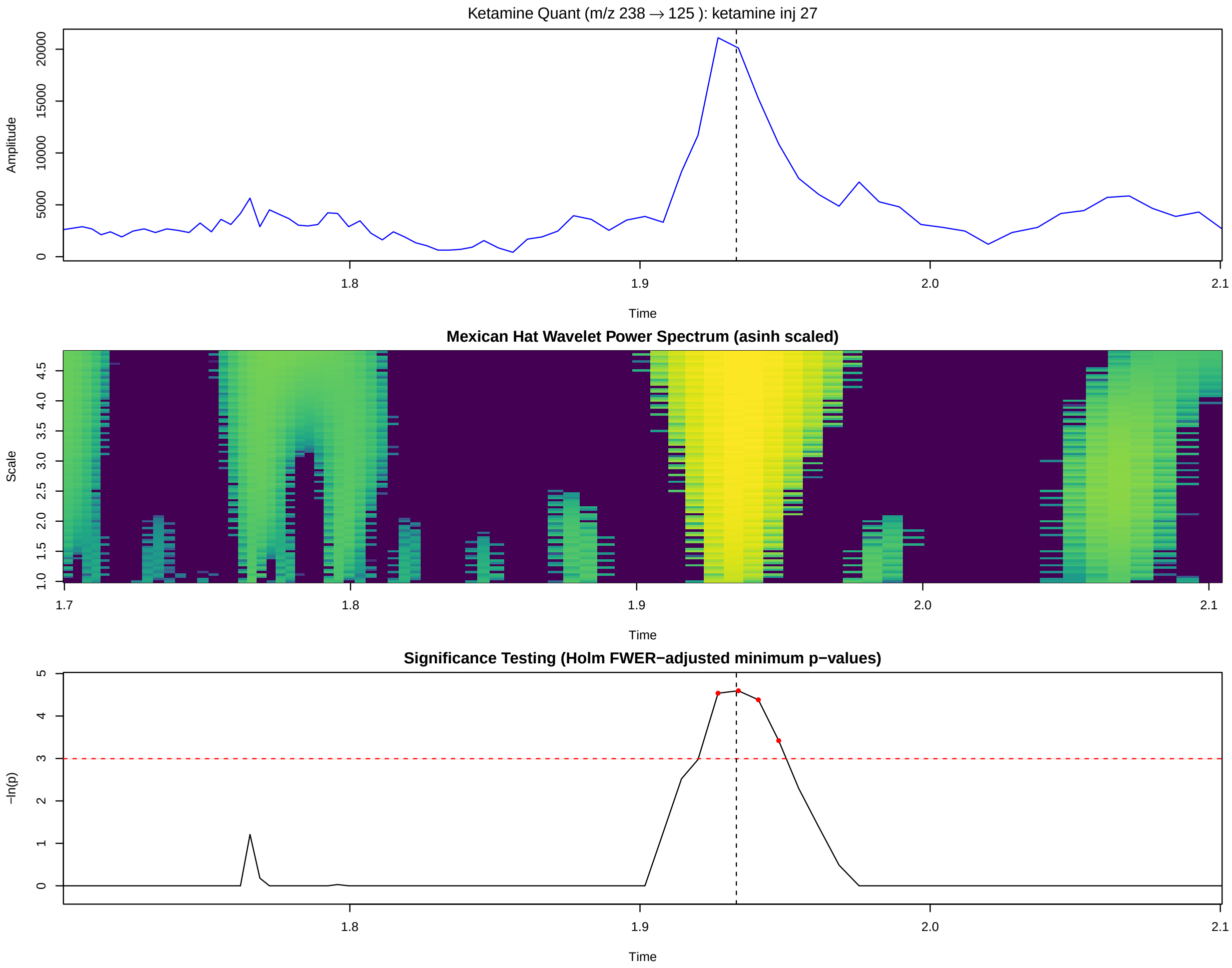


Figure S3. Ketamine sub-LLOQ biological positive (Inj 27) significance test on the SCIEX 6500. Quantifier transition m/z 238 → 125. This confirmed positive sample has signal intensity below the low calibration standard, yet the wavelet significance test detects 4 significant time points ($p_{\mathrm{adj}} = 1.01 \times 10^{-2}$) at the expected retention time.

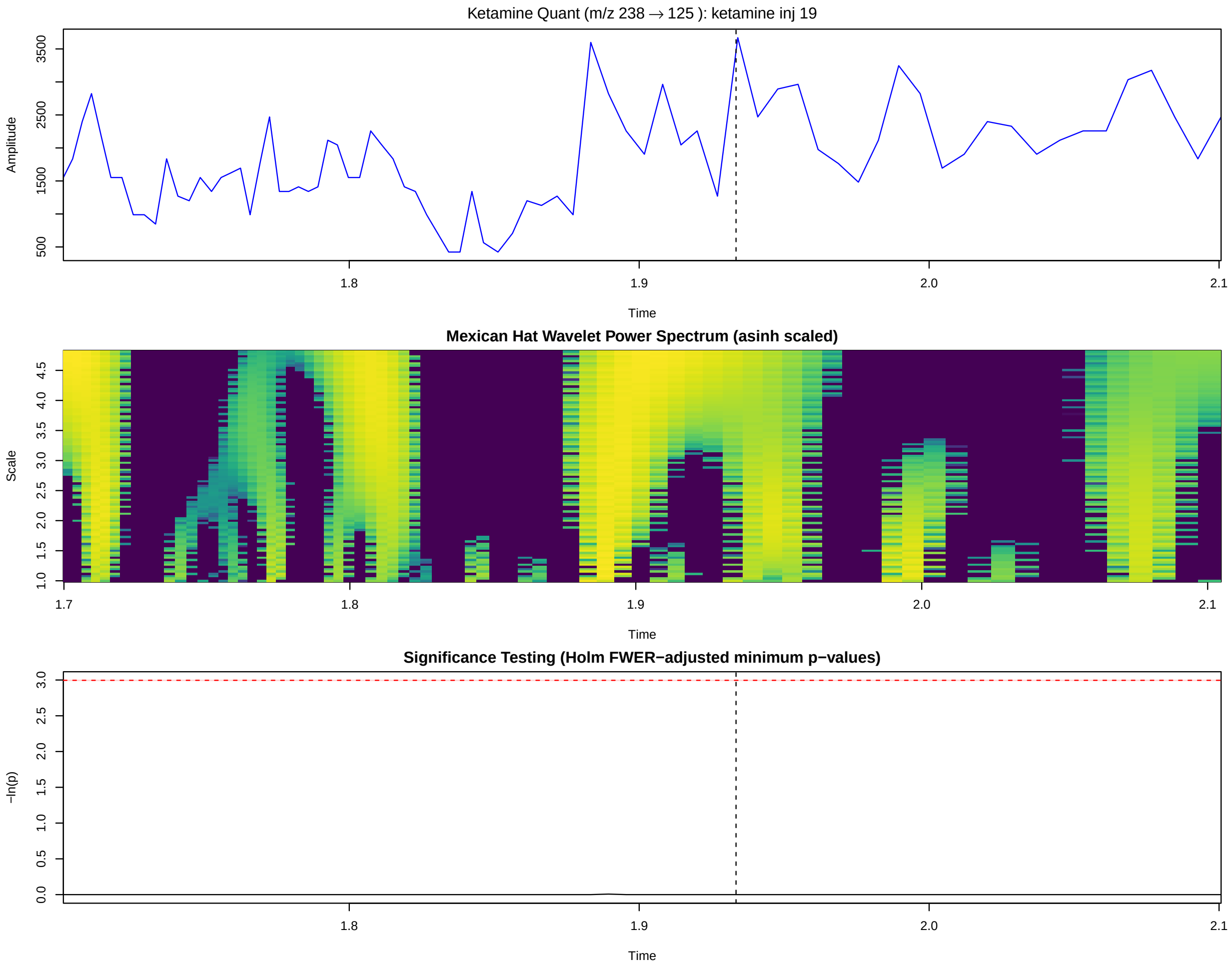


Figure S4. Ketamine biological negative (Inj 19) significance test on the SCIEX 6500. Quantifier transition m/z 238 → 125. No significant features detected ($p_{\text{adj}} = 0.993$). The chromatographic features in this sample are visually similar to peaks that might be manually integrated in conventional analysis, but the Monte Carlo significance test correctly classifies them as consistent with the chemical noise null model.

Per-sample chromatograms, wavelet power maps, chemical noise characterizations, and significance test results for all 16 ketamine samples are available in the project repository at https://github.com/rkjulian/wavelet-peak-significance.

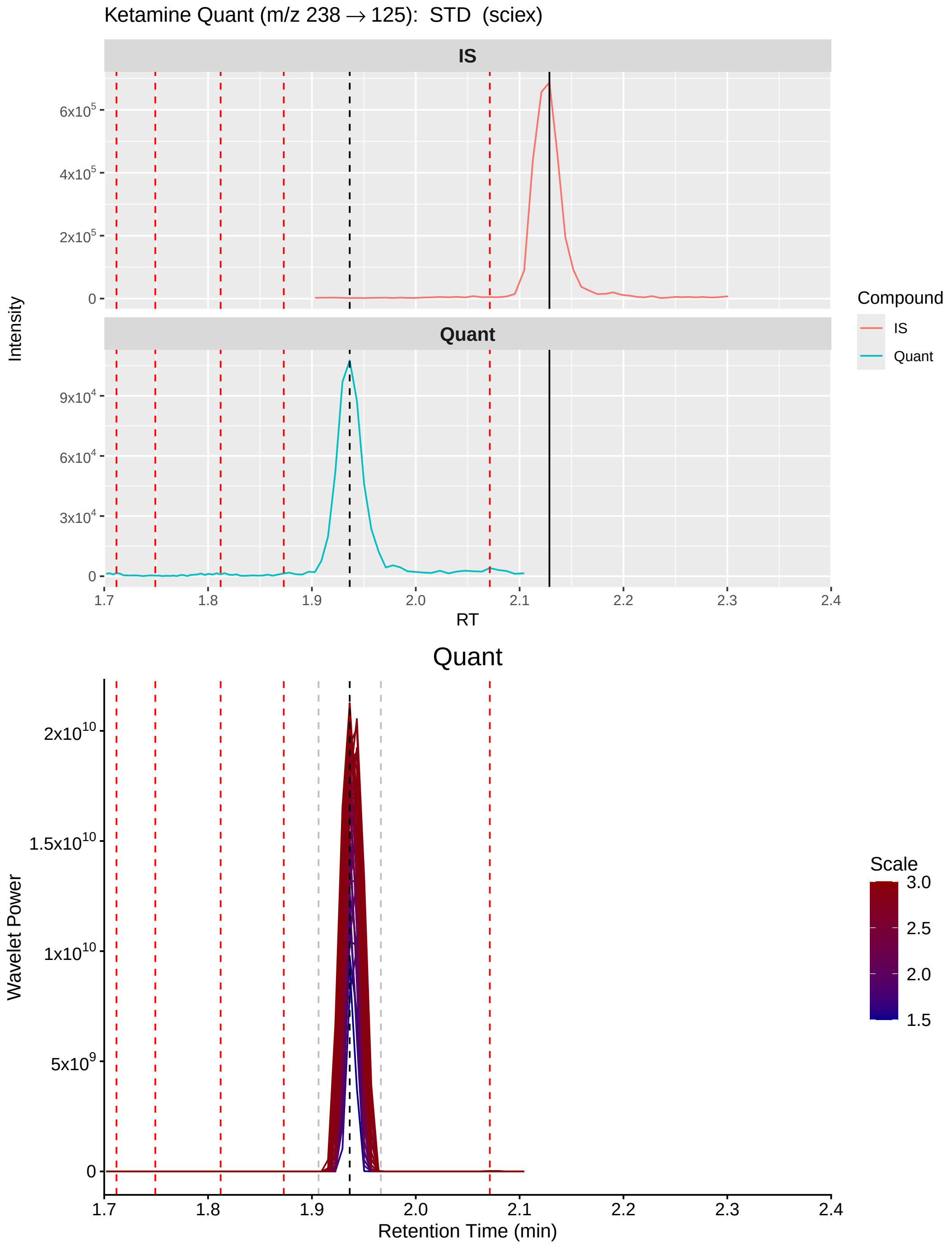


Figure S5. Ketamine chemical noise characterization for the low standard (STD) on the SCIEX 6500. Quantifier transition m/z 238 $\rightarrow$ 125. Top panels: IS and Quant chromatograms with detected chemical noise peaks (red dashed lines) and expected quant retention time (black dashed line). Bottom: wavelet power spectrum across scales 1.5–3.0 with quant RT proximity bounds (gray dashed lines).

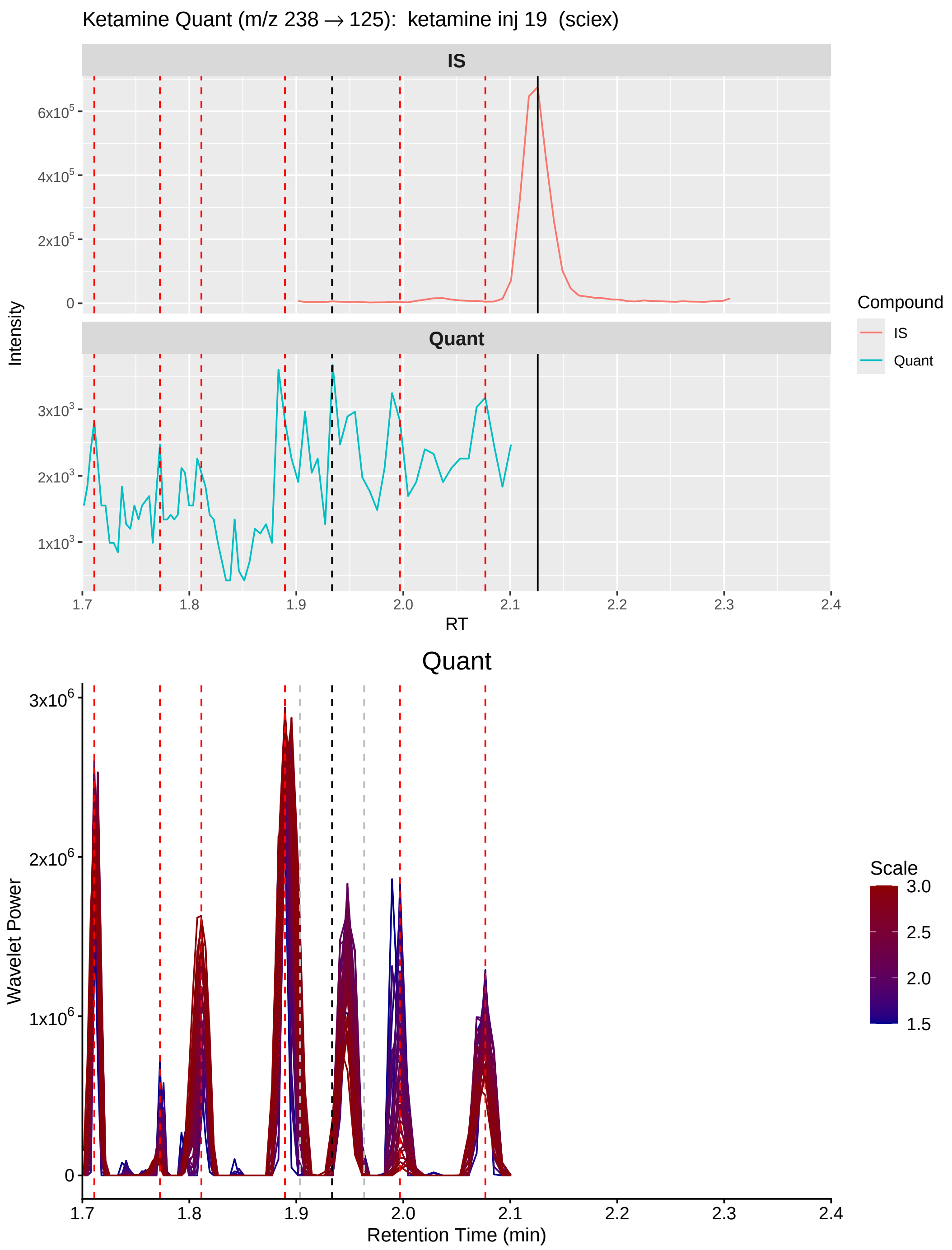


Figure S6. Ketamine chemical noise characterization for Inj 19 on the SCIEX 6500. Quantifier transition m/z 238 → 125. The wavelet power spectrum illustrates the level of structured chemical background in a biological negative sample. Chromatographic features visible in the raw trace do not produce wavelet power exceeding the chemical noise null model, consistent with the non-significant result in the significance test (Figure S4).
